# Systemic *in vivo* phage selection reveals compartment-dependent organization of recoverable peptide repertoires

**DOI:** 10.64898/2026.04.16.719084

**Authors:** Junko Okano, Miwako Katagi, Kazunori Fujino, Atsunori Shindo, Yoshio Furusho, Hideto Kojima

## Abstract

Previous *in vivo* phage display studies establish the value of the approach for identifying organ-homing and tissue-associated peptides, but whether recovered sequences form diffuse hit lists or structured repertoires remains unclear. This study combines systemic *in vivo* phage selection, laser-capture microdissection, and high-throughput sequencing to profile recoverable 7-mer peptide repertoires across 41 bulk organs, 39 capillary beds, and 125 parenchymal microenvironments. Across these anatomical scales, the repertoires show non-diffuse, compartment-dependent organization in enrichment-profile space, supported by robustness and null-model analyses. Compartment resolution provides information not apparent in bulk-organ profiles: representative motif-defined neighborhoods exhibit contrasting local sequence-tolerance regimes, ranging from bounded variation to near-single-sequence convergence. Compartment-resolved analysis also nominates EVGTARY, a cortex-associated candidate that is not prioritized by bulk-organ ranking. In an exploratory mouse-level analysis, an EVGTARY-conjugated branched polyethyleneimine (BPEI) polyplex is associated with increased cortical DsRed fluorescence compared with BPEI polyplex alone. Together, these findings shift the analytical unit of *in vivo* phage display from isolated organ-associated hits toward the organization of recoverable peptide repertoires and support an interpretation in which intact tissue context constrains the range of peptide solutions that remain recoverable after systemic *in vivo* phage selection.

## 1. Introduction

Tissue-selective peptide discovery has focused on identifying individual sequences that are recovered from, or enriched in, a target organ (*1,2*). However, systemically administered phage-displayed peptide libraries encounter a series of biological contexts before recovery, including vascular transport, tissue exposure, microanatomical organization, and local molecular environments. These processes may influence not only which peptides are recovered, but also how the recoverable repertoire is organized across tissues. Therefore, a central unresolved question is whether systemic *in vivo* phage selection yields a collection of largely isolated tissue-associated hits or a structured peptide repertoire whose organization varies across anatomical contexts.

*In vivo* phage display provides an experimental means of sampling peptide recovery directly after systemic circulation through intact organisms. Since the initial identification of organ-associated homing peptides, this approach has revealed vascular heterogeneity and organ-associated molecular addresses, including so-called vascular zip codes (3,4). Subsequent studies have identified tumor-homing and tissue-penetrating peptides and have extended vascular peptide mapping to humans (5–7). More recently, high-throughput sequencing has enabled increasingly quantitative analysis of phage-displayed peptide libraries and selected pools (8–11). Despite these advances, most *in vivo* phage-display studies have continued to emphasize whole-organ enrichment profiles or individual candidate sequences. Whole-organ measurements integrate vascular, stromal, and parenchymal contributions into a single composite readout, making it difficult to determine how peptide recovery is organized across tissue compartments or whether compartment-resolved profiling provides information that is qualitatively different from bulk-organ ranking.

Here, we combined systemic multi-organ *in vivo* phage selection with laser-capture microdissection and next-generation sequencing to profile recoverable peptide repertoires across 41 whole organs, 39 capillary beds, and 125 parenchymal microenvironments. By treating recovered peptides as high-dimensional repertoire data, we examined whether their enrichment profiles formed structured organization across anatomical scales and whether compartment resolution provided information distinct from whole-organ analysis. The resulting landscapes revealed organ- and compartment-associated repertoire organization, distinct local motif-defined sequence-tolerance regimes, and a compartment-nominated candidate that was not prioritized by bulk-organ ranking. Together, these findings support a repertoire-level view of systemic *in vivo* phage selection and an interpretation of tissue-constrained peptide recovery.

## 2. Results

### 2.1 Global embeddings reveal structured repertoire organization across anatomical scales

As illustrated in Fig. 1A, *in vivo* phage display was performed across 41 bulk organs. To resolve tissue compartments, selected tissues were further microdissected into 39 capillary beds and 125 parenchymal microenvironments using laser-capture microdissection (LCM). We treated each recovered peptide as a sample-wise enrichment profile and used unsupervised Uniform Manifold Approximation and Projection (UMAP) to visualize the global organization of the bulk, capillary, and parenchymal repertoires. Across the three datasets, the embeddings showed shared central regions together with peripheral structures associated with physiological systems or individual organs, while exhibiting distinct overall geometries at each anatomical scale. The dimensional properties, robustness, and null-model behavior of these embeddings were evaluated in supplementary analyses (figs. S1–S7).

**Fig. 1.**
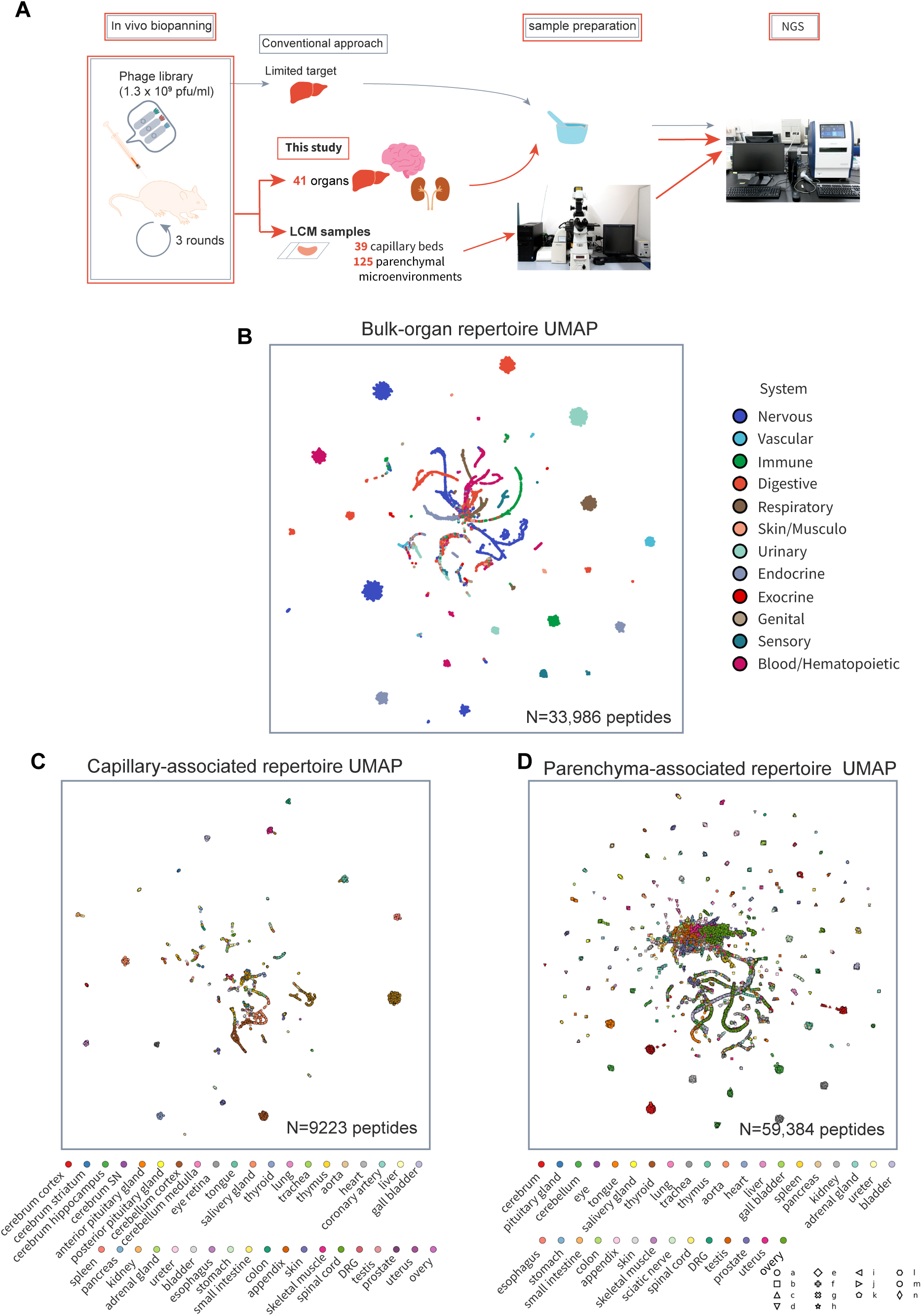
Systematic mapping of *in vivo*-selected recoverable peptide repertoires across whole-organ and compartment-resolved scales. **A) Schematic overview of the study design.** The conventional workflow shown in gray represents *in vivo* biopanning focused on a limited number of target organs, whereas the present study, shown in orange, combined systemic *in vivo* phage selection across multiple organs with whole-organ and laser-capture microdissection (LCM)-based compartment-resolved profiling. Phage-derived sequences recovered from 41 bulk organs, 39 capillary beds, and 125 parenchymal microenvironments were profiled by next-generation sequencing and analyzed at the repertoire level. The final available selected sample corresponded to round 2 for the urinary bladder and round 3 for all other organs. **B) UMAP embedding of peptide enrichment profiles across 41 bulk organs.** Each point represents a unique recovered 7-mer peptide and is colored according to its physiological system annotation. Detailed organ-level overlays are provided in fig. S4, and the system–organ mapping is listed in table S1. **C) UMAP embedding of peptide enrichment profiles across 39 LCM-derived capillary beds.** Each point represents a unique recovered 7-mer peptide and is colored according to the associated organ. **D) UMAP embedding of peptide enrichment profiles across 125 LCM-derived parenchymal microenvironments.** Each point represents a unique recovered 7-mer peptide and is colored according to the associated organ. Letters a–n indicate anatomical subregions listed in table S2. The number of peptides included in each embedding is indicated in panels (B–D). UMAP proximity reflects similarity in organ-wise or sample-wise enrichment profiles rather than direct amino-acid sequence similarity. Bulk-organ profiles integrate vascular and parenchymal contributions into a composite readout and therefore do not resolve the compartmental origin of organ-associated enrichment. Embedding robustness, parameter-sensitivity analyses, and null-model assessments are provided in figs. S1–S7. Abbreviations: DRG, dorsal root ganglion; SN, substantia nigra.

In the bulk-organ embedding (Fig. 1B), a shared central region gave rise to radially extending system-associated structures, while multiple peripheral islands were enriched for individual physiological systems. Detailed system–organ annotations are provided in table S1. The overall organization was reproducible across alternative UMAP initializations and clustering-parameter settings (figs. S2–S5). In contrast, permutation of the system labels dispersed the apparent system-level color coherence, whereas feature-wise shuffling of the peptide-by-organ matrix disrupted the shared central organization and increased global spread (fig. S6). These results indicate that the bulk-organ embedding reflects reproducible organization in the original peptide–organ enrichment profiles rather than visualization instability or random peptide–organ correspondence.

In the capillary-associated embedding (Fig. 1C), peptides derived from multiple organs formed a shared central region surrounded by a largely continuous and organ-mixed structure. Short organ-associated extensions were present, but the global organization remained comparatively overlapping, with isolated organ-enriched islands distributed at the periphery. Silhouette analysis was consistent with a more continuous and less separable organization in the capillary dataset than in the bulk-organ and parenchymal datasets (fig. S7).

In the parenchyma-associated embedding (Fig. 1D), a compact shared region was accompanied by elongated organ-associated structures extending from the center. Several of these structures contained multiple anatomically related subregions arranged along continuous paths, indicating that within-organ anatomical variation contributed to the organization of the parenchymal repertoire. Additional organ-enriched clusters and isolated peripheral islands were distributed throughout the embedding; organ and subregion annotations are provided in table S2.

Together, these embeddings provide an exploratory map of structured organ- and compartment-associated repertoire organization in enrichment-profile space. However, the embedding geometry alone does not quantify organ-restricted enrichment from broadly shared enrichment across organs. Therefore, we investigated organ-level enrichment patterns in the bulk *in vivo* biopanning data using a digital-subtraction analysis.

### 2.2 Multi-organ digital subtraction identifies a cerebrum-restricted upper-ranking regime

To quantify organ-restricted endpoint enrichment in the bulk-organ dataset, we calculated peptide enrichment from round 1 and the final available selected-round read counts and defined a digital subtraction score as enrichment in the cerebrum minus the maximum enrichment observed across all other organs. The cerebrum was selected as a focused example because it showed strong separation in the bulk-organ embedding (fig. S4). Ranking peptides by this score revealed a cerebrum-restricted upper-ranking regime (Fig. 2A).

**Fig. 2.**
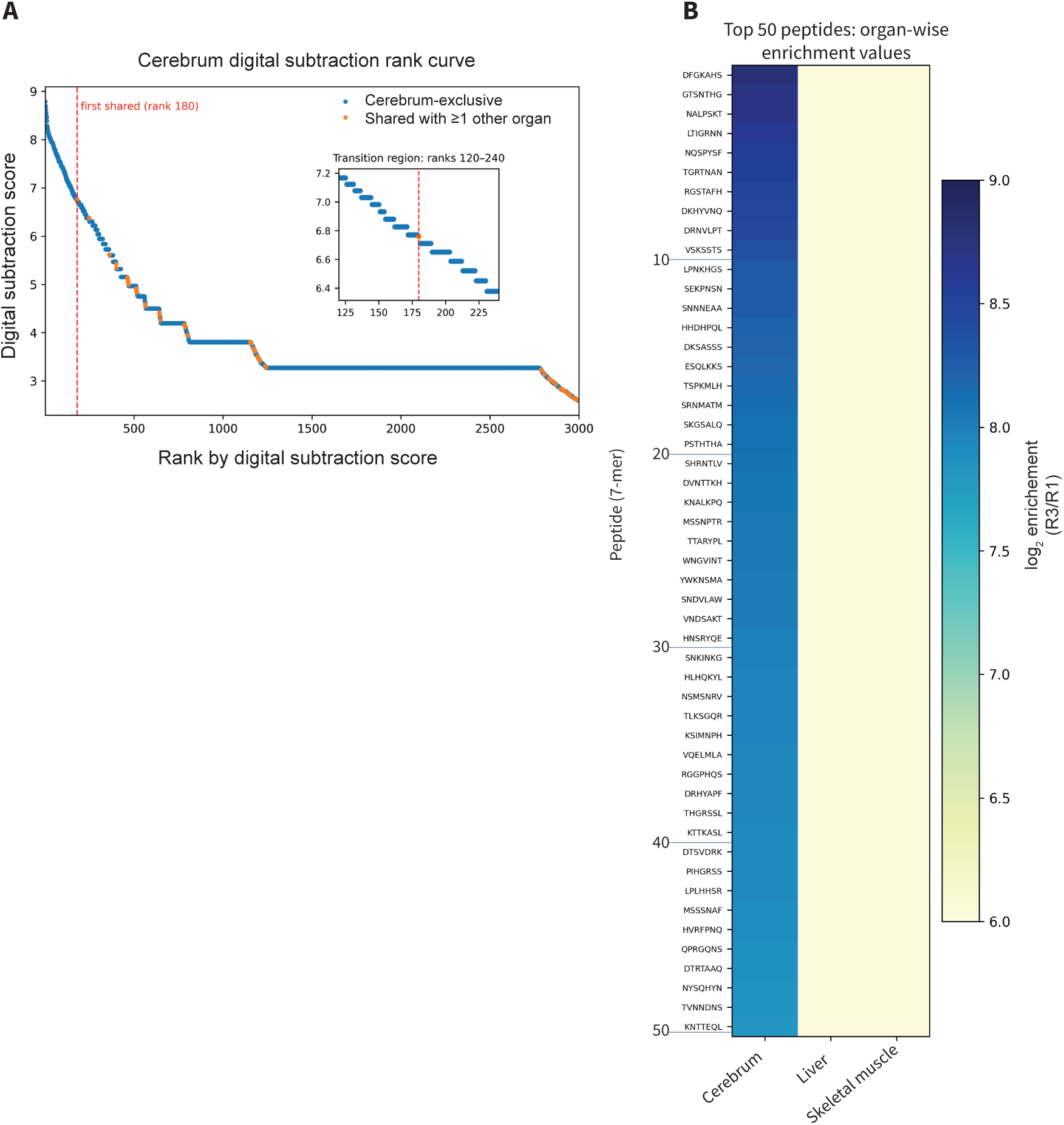
Multi-organ digital subtraction identifies a cerebrum-restricted upper-ranking regime in bulk data. **A) Cerebrum digital subtraction rank curve derived from the bulk-organ dataset.** Peptides were ranked using a digital subtraction score defined as enrichment in the cerebrum minus the maximum enrichment observed across all other organs. Cerebrum-exclusive peptides, defined as peptides with positive enrichment only in the cerebrum, are shown in blue, whereas peptides with positive enrichment in at least one additional organ are shown in orange. The first shared peptide occurred at rank 180, marked by the vertical dashed line, indicating a transition from a cerebrum-exclusive upper-ranking regime to a shared regime. The inset shows the transition region spanning ranks 120–240. **B) Organ-wise enrichment of the top 50 peptides ranked by the cerebrum digital subtraction score.** Enrichment values are shown for the cerebrum, liver, and skeletal muscle. The top-ranked peptides showed strong enrichment in the cerebrum, whereas enrichment in the liver and skeletal muscle remained near background levels, confirming that the upper-ranking regime was dominated by cerebrum-restricted enrichment in the bulk-organ dataset.

Specifically, the top 179 ranked sequences were enriched exclusively in the cerebrum, and the first peptide showing enrichment in additional organs (“shared”) appeared at rank 180 (Fig. 2A and its inset). The plateau region of the rank curve consisted entirely of cerebrum-exclusive peptides, whereas shared peptides were predominantly distributed in regions in which the subtraction score decreased progressively. Examination of the full-rank distribution (ranks 1–30,000) further showed that the subtraction score did not decrease smoothly or monotonically from top to bottom. Instead, the rank curve exhibited a stepwise decline with multiple plateau regions, suggesting a hierarchical structure in the ranked distribution (fig. S8).

Consistent with the rank curve, heatmap analysis of representative organs showed that the top-ranked sequences were strongly enriched in the cerebrum, whereas enrichment in the liver and skeletal muscle remained near the background levels (Fig. 2B). These findings confirm that, at least within the upper-ranking regime, high subtraction scores are driven primarily by strong cerebrum-specific enrichment. Detailed enrichment values for the representative top-ranked peptides are provided in table S3.

However, because the bulk dataset represents a mixture of multiple tissue components, it remains unclear whether the observed enrichment arises from the vascular compartment or from the parenchymal tissue. This limitation led us to analyze the compartment-resolved capillary and parenchymal datasets to examine whether the selected peptide neighborhoods were shaped by distinct compartment-level constraints.

### 2.3 Motif analysis reveals local sequence-constraint regimes

To dissect local sequence constraints within representative compartment-associated neighborhoods, we performed motif-centered analyses anchored by medoid sequences extracted from the capillary and parenchymal embedding spaces. Cross-compartment overlap analysis showed strong overlap between the capillary and parenchymal datasets, with an observed-to-expected enrichment of 124.6 and a nominal one-sided hypergeometric *P* value of 1.3 × 10^-222^, motivating focused analysis of the shared peptide structure across these two compartments.

Distinct cluster-central medoid sequences were identified in the two datasets: STLHQEL in the capillary dataset and STFHQKL in the parenchymal dataset (Fig. 3A). Consistent with their embedding-based assignment, organ-level heatmaps based on log1p-transformed raw read counts confirmed compartment-associated enrichment of these medoid sequences (fig. S9). The two medoids also occupied nearby regions along a shared embedding axis, indicating local continuity between the capillary- and parenchyma-associated enrichment-profile landscapes (fig. S10).

**Fig. 3.**
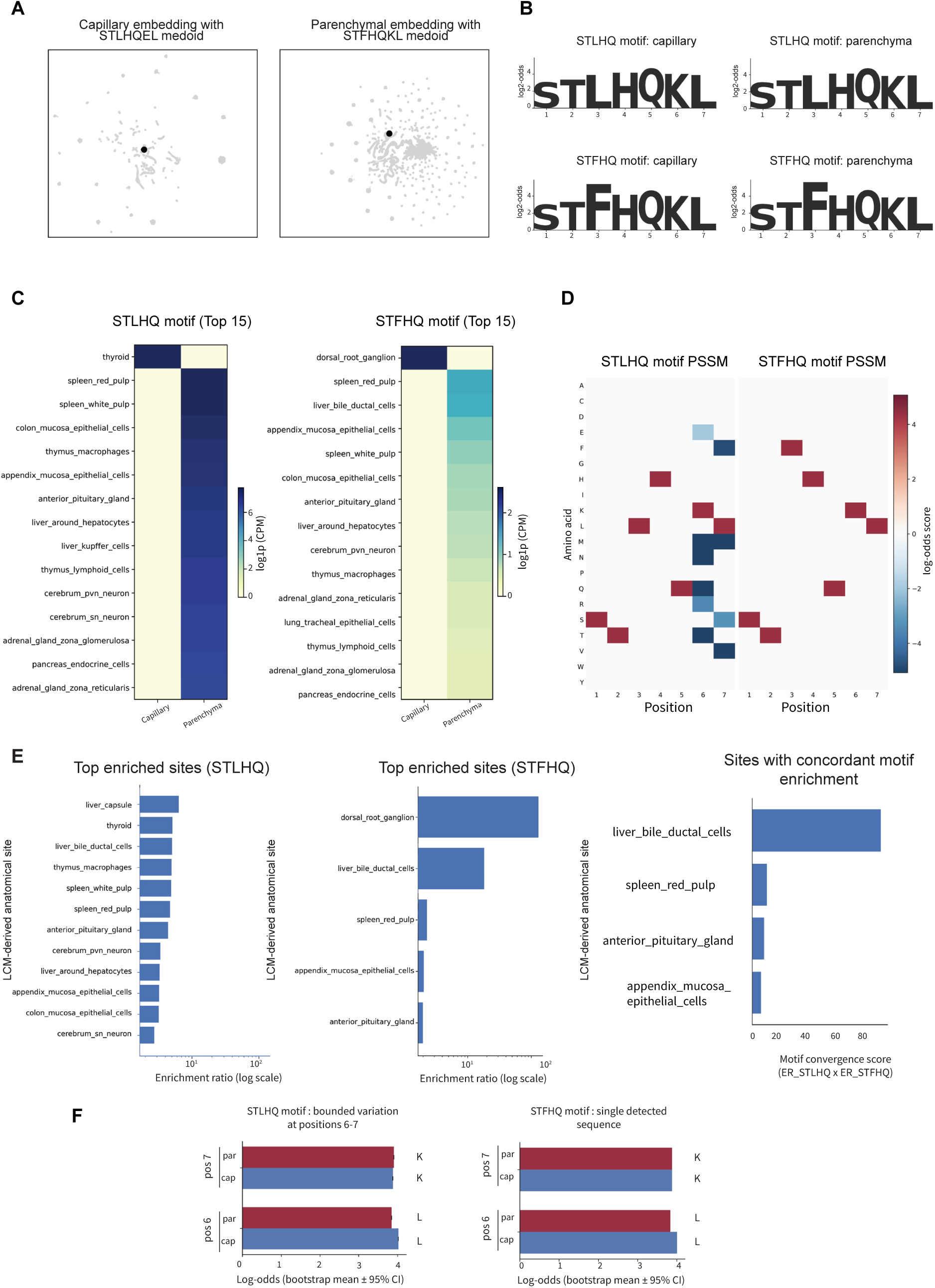
Representative motif-defined peptide neighborhoods exhibit distinct local sequence-constraint regimes. **A) Geometry-derived medoid sequences in the capillary and parenchymal embeddings.** STLHQEL and STFHQKL (black points) were identified as cluster-central medoids in the capillary-associated (left) and parenchyma-associated (right) embedding spaces, respectively. Supporting analyses of medoid enrichment and embedding relationships are shown in figs. S9 and S10. **B) Read-weighted sequence logos of motif-defined peptide sets anchored by the capillary and parenchymal medoids.** Positive log₂-odds scores are shown for the Tier 0 motifs (^STLHQ..$ and ^STFHQ..$) in each compartment. Because the logos summarize the read-weighted composition of the complete motif-defined peptide sets, their dominant consensus sequences do not necessarily coincide with the geometry-derived medoids shown in panel (A). **C) Abundance of STLHQ- and STFHQ-matching peptide sets across the top 15 LCM-derived anatomical sites.** Heatmaps show log1p-transformed CPM values in the capillary and parenchymal datasets. Color scales were independently normalized within each motif-defined peptide set. Heatmaps including all anatomical sites are shown in fig. S12. **D) Full position-specific scoring matrix (PSSM) heatmaps for the STLHQ- and STFHQ-defined peptide sets.** Zero-centered color scaling shows position-specific amino-acid enrichment and depletion relative to a uniform 20-amino-acid background. **E) Anatomical-site enrichment and concordance across the two motif-defined peptide sets.** Left and middle, enrichment ratios are shown for significantly enriched LCM-derived anatomical sites for the STLHQ- and STFHQ-defined peptide sets, respectively (q < 0.05). For STLHQ, a representative subset of significantly enriched sites is displayed; the complete set is shown in fig. S14. For STFHQ, all sites meeting the significance threshold (n = 5) are shown. Right, sites meeting the significance criterion in both motif-defined analyses are ranked using a motif convergence score calculated as the product of the corresponding enrichment ratios. **F) Bootstrap-based assessment of positional constraint within the motif-defined peptide sets.** Left, bootstrap means and 95% percentile intervals are shown for log₂-odds scores at positions 6 and 7 of the STLHQ-defined peptide sets in the capillary and parenchymal datasets. Right, the corresponding estimates are shown for the STFHQ-defined peptide sets. No visible intervals are present for STFHQ because the bootstrap distributions collapsed to the point estimates, consistent with single-sequence representation in both compartments. Abbreviations: cap, capillary; ER, enrichment ratio; par, parenchyma; pos, position.

To characterize local sequence variation around each medoid, Tier 0 motifs (^STLHQ..$ and ^STFHQ..$) were analyzed in both datasets. Sequence composition was visualized using log-odds-based sequence logos (Fig. 3B). Because positions 1–5 were fixed by the motif definitions, the logos were used primarily to visualize read-weighted amino-acid usage at the variable positions 6 and 7 across compartments. These descriptive profiles motivated subsequent quantitative assessment of positional tolerance.

Because the capillary and parenchymal datasets differed substantially in size, motif abundance was compared using log1p-transformed CPM values. Empirical cumulative distribution function analysis showed comparable distributional shapes between the datasets despite a global shift in abundance (fig. S11), supporting normalized, distribution-aware comparison between compartments. On this basis, the motif-associated abundance was ranked separately for the STLHQ- and STFHQ-defined sets within each compartment, and the top 15 LCM-derived anatomical sites were visualized as heatmaps (Fig. 3C). For both motifs, the highest-ranking sites were distributed across capillary and parenchymal samples rather than confined to the compartment from which the medoid was derived, indicating that motif-associated signals extended across tissue compartments. Heatmaps including all anatomical sites are shown in fig. S12. Comparable contrast patterns were also observed when raw read counts were directly visualized (fig. S13).

To further quantify positional constraints, full position-specific scoring matrices (PSSMs) were constructed for the Tier 0 motifs after combining compartments and were visualized as heatmaps (Fig. 3D). For the STLHQ motif, strong negative scores were concentrated primarily at positions 6 and 7, indicating restricted substitution tolerance at these sites. In contrast, the STFHQ motif showed strong positive scores for the consensus residues across positions 1–7, consistent with markedly stronger convergence toward a dominant sequence. Together, these analyses indicated that the two motif-defined neighborhoods differed not only in their enrichment patterns but also in the degree of tolerated sequence variation.

To account for differences in total read abundance across LCM-derived anatomical sites, enrichment ratios were calculated for each motif-defined set at each site (Fig. 3E). Multiple sites showed significant enrichment of the STLHQ-defined set, whereas significantly enriched sites for the STFHQ-defined set were fewer and more restricted (Fig. 3E, left and middle; fig. S14). To identify sites exhibiting concordant enrichment across both motif-defined sets, we defined a motif convergence score as the product of the corresponding enrichment ratios. Among sites showing significant enrichment in both analyses (q < 0.05 for each component analysis), liver bile ductal cells had the highest motif convergence score, followed by spleen red pulp, anterior pituitary gland, and appendix mucosa epithelial cells (Fig. 3E, right). The individual enrichment ratios underlying these scores are shown in fig. S15, and the distribution of statistical significance across all sites is shown in fig. S16, with corresponding statistics provided in table S4.

We next evaluated motif stability and tolerance boundaries using read-weighted sequence-level bootstrap resampling and tiered perturbation analyses (Fig. 3F and fig. S17). Bootstrap statistics were computed for all motif positions, but interpretation focused on positions 6 and 7, the variable positions permitted under the Tier 0 motif definitions. For the STLHQ-defined set, these positions showed narrowly bounded estimates across resampling (Fig. 3F, left). In contrast, the STFHQ-defined set showed no visible bootstrap intervals at positions 6 and 7, consistent with representation by a single detected sequence in both compartments (Fig. 3F, right).

Under progressively relaxed pattern definitions (Tier 3–Tier 4), rare low-frequency deviations became detectable at selected positions (fig. S17), yet the consensus residues remained overwhelmingly dominant. The variant counts remained limited across the Tier 0–Tier 4 conditions (table S5), indicating that the STLHQ-defined neighborhood retained a bounded local solution space within the observed repertoire. In contrast, the STFHQ-defined set remained represented by a single detected sequence in both compartments. No alternative variants emerged across the tested relaxed motif definitions, consistent with near-complete convergence within the observed repertoire.

To assess whether the observed differences in sequence diversity between the capillary and parenchyma-associated STLHQ peptide sets could be explained by read-depth differences alone, the read-weighted Shannon entropy was calculated across tiers (fig. S18). The entropy values remained stable within each compartment across progressive relaxation of the motif constraints, with the parenchyma-associated STLHQ peptide set consistently showing higher entropy than the capillary-associated set. Therefore, within the STLHQ-defined peptide sets, these results are consistent with greater tolerated sequence diversity in the parenchyma-associated subset than in the capillary-associated subset.

Collectively, these analyses show that the representative STLHQ-defined neighborhood was sharply bounded yet retained limited sequence variation, whereas the STFHQ-defined neighborhood showed pronounced convergence toward a single detected sequence. Despite these distinct local sequence-constraint regimes, both motif-defined sets showed concordant enrichment across a restricted subset of LCM-derived anatomical sites (Fig. 3E, right). However, whether a peptide nominated from these compartment-resolved landscapes could yield a detectable *in vivo* signal in an initial proof-of-concept test remained unresolved.

### 2.4 Compartment-resolved landscape analysis nominates EVGTARY for exploratory *in vivo* evaluation

As an exploratory test of candidate nomination from compartment-resolved landscapes, we selected EVGTARY from the cortex-associated parenchymal landscape for *in vivo* evaluation. In the global embeddings of the capillary and parenchymal datasets, EVGTARY was located away from the respective medoids, STLHQEL and STFHQKL (Fig. 4A, left). However, when peptides associated with brain-related enrichment were extracted and reclustered using hierarchical density-based spatial clustering of applications with noise (HDBSCAN), EVGTARY localized near the medoid of the corresponding local cluster (Fig. 4A, right). EVGTARY was detected almost exclusively in cortex-associated parenchymal samples and was not detected in other parenchymal organs (fig. S19). In contrast, it was not among the top-ranked peptides in the bulk cerebrum dataset (table S3).

**Fig. 4.**
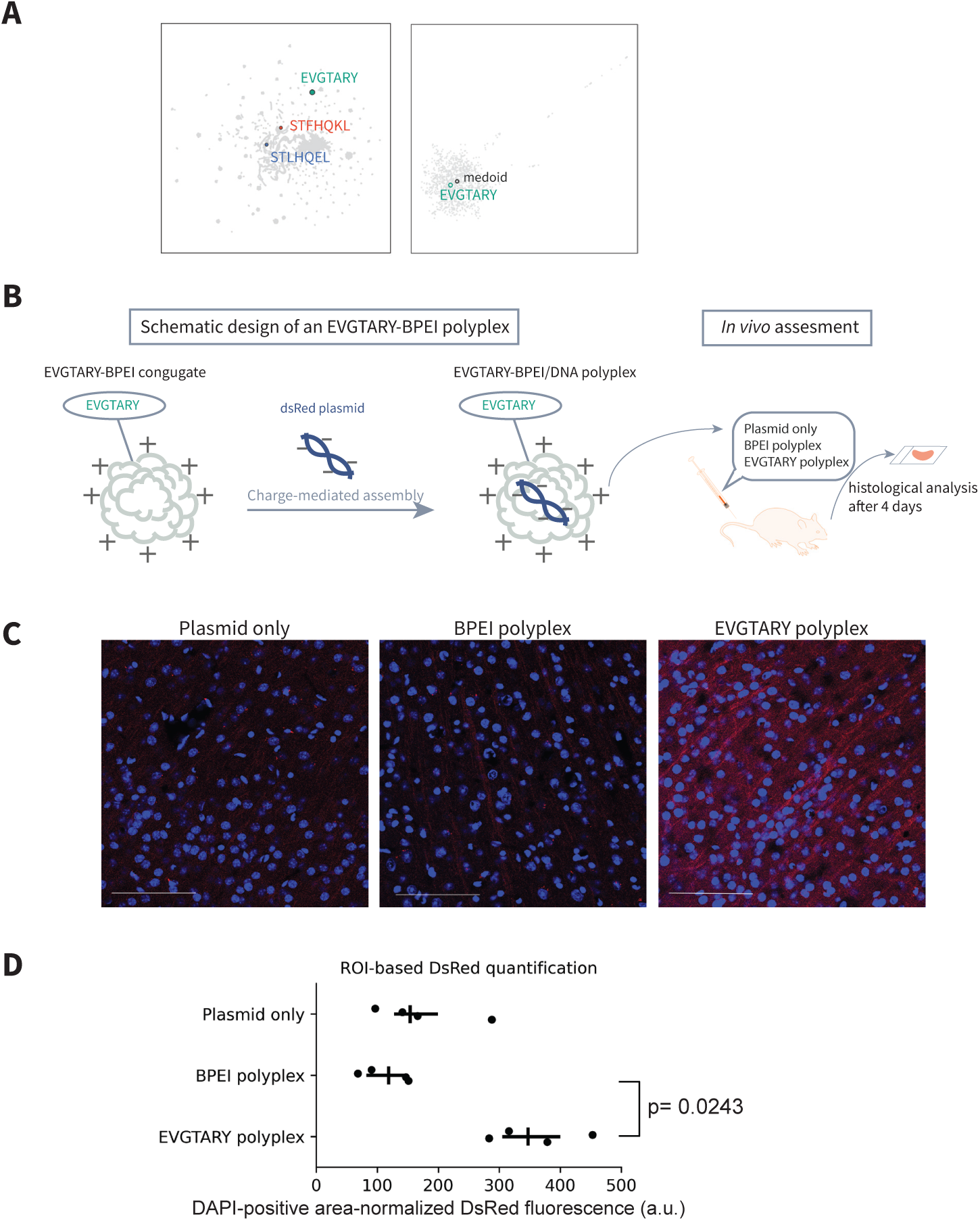
Exploratory in vivo evaluation of a parenchymal-landscape-derived brain-associated candidate. **A) Selection of EVGTARY from LCM-derived peptide embeddings.** Left, EVGTARY (green) is shown within a unified embedding of capillary and parenchymal peptide datasets together with representative medoid sequences, STLHQEL (blue) and STFHQKL (orange). Right, in the brain-associated subset reclustered by using HDBSCAN, EVGTARY localizes near the cluster medoid (black), with a medoid distance of 0.1237 (ranked 95th of 1,098 sequences; 8.65th percentile). **B) Schematic illustration of the proof-of-concept design and *in vivo* evaluation.** An EVGTARY–branched polyethyleneimine (BPEI) polyplex was assembled and systemically administered. Control groups included dsRed plasmid alone (plasmid only) and a BPEI/DNA polyplex lacking the EVGTARY peptide (BPEI polyplex). Histological analyses were performed 4 days after systemic administration. **C) Representative cortical sections are shown for the plasmid-only, BPEI-polyplex, and EVGTARY-polyplex groups.** Stronger DsRed fluorescence was visually observed in the representative EVGTARY-polyplex section. Images were acquired under identical microscope settings and displayed with identical contrast and lookup table (LUT) scaling across all conditions. Scale bars, 100 µm. **D) ROI-based quantification of DsRed fluorescence.** DsRed integrated density was normalized to the DAPI-positive area within the same ROI. Each point represents one mouse; two microscopic fields were averaged per mouse (n = 4 mice per group). Group differences were assessed by Kruskal–Wallis test followed by Dunn’s post hoc test with Benjamini–Hochberg correction. EVGTARY polyplex increased DAPI-positive area-normalized DsRed fluorescence compared with BPEI polyplex alone (adjusted P = 0.0243). Abbreviation: a.u., arbitrary units.

To assess whether EVGTARY and the representative motif features identified after *in vivo* selection were already prominent in the input library, we performed same-lot naïve-library background profiling using L35742 and L35743, two sequencing profiles generated from the same unamplified naïve phage library lot used in the mouse experiments. The exact sequences STLHQEL, STFHQKL, and EVGTARY were not detected in either profile. Likewise, no valid 7-mer matching the ^STLHQ..$ or ^STFHQ..$ motif was detected in either dataset (fig. S20). Thus, these specific selected features were not detected in the sequenced same-lot input-library profiles. Because the naïve-library data were not incorporated into downstream normalization or landscape analyses, they were used for input-background profiling rather than direct naïve normalization.

To assess EVGTARY *in vivo*, the peptide was conjugated to branched polyethyleneimine (BPEI), and EVGTARY–BPEI/DNA polyplexes were assembled by charge-mediated complex formation with a dsRed plasmid (Fig. 4B). The BPEI-to-DNA ratios used for polyplex formation were determined using a gel-retardation assay (fig. S21). Polyplexes corresponding to 2.04 μg of DNA were administered by tail-vein injection, and tissues were analyzed histologically four days after administration.

Representative cortical sections showed stronger DsRed fluorescence in the EVGTARY-polyplex group than in the plasmid-only and BPEI-polyplex groups (Fig. 4C). Quantitative group comparisons were then performed by normalizing DsRed integrated density to the DAPI-positive area within the same cortical ROI. Two microscopic fields were averaged within each mouse before statistical analysis. As a result, there was an overall difference among the three treatment groups (Kruskal–Wallis test, *P* = 0.0264; Fig. 4D). Dunn’s post hoc test with Benjamini–Hochberg correction showed that the EVGTARY polyplex significantly increased DAPI-positive area-normalized DsRed fluorescence compared with BPEI polyplex alone (adjusted *P* = 0.0243).

In contrast, in the liver, where background fluorescence was relatively high, the signals remained at background levels across all groups, and almost no signal was detected in the small intestine (fig. S22).

Together, these results provide a proof-of-concept observation that an EVGTARY-conjugated BPEI polyplex is associated with increased cortical DsRed signal after systemic administration, with a significant increase over the BPEI polyplex alone.

## 3. Discussion

Systemic *in vivo* phage selection has often been interpreted through the recovery of individual organ-associated peptide hits (*3,6,12,13*). Our findings extend this analytical scope by showing that recoverable peptides can also be examined as structured repertoires across anatomical scales. By treating *in vivo*-selected peptides as high-dimensional repertoire data and adapting unsupervised embedding and clustering strategies commonly used in single-cell analysis to peptide enrichment profiles, we identified non-diffuse, compartment-dependent organization across whole-organ, capillary, and parenchymal datasets. These analyses do not represent direct maps of peptide–receptor binding; rather, they describe how peptide recovery is organized after systemic circulation, tissue exposure, selection, and recovery. The convergence of global repertoire organization with organ-restricted enrichment and local motif-defined sequence restriction supports the interpretation that intact tissue context limits the range of peptide solutions that remain recoverable after systemic *in vivo* phage selection.

Previous *in vivo* phage-display studies have established the value of the approach for identifying vascular addresses, organ-homing ligands, and tissue-penetrating candidates (*3–7,10,12,13*). Although high-throughput and compartment-oriented variants have emerged (*9,11,12*), the field has largely remained focused on identifying organ- or disease-associated candidates rather than analyzing body-wide repertoire organization across matched anatomical compartments. In the present study, a shared central region and peripheral system- or organ-associated structures demonstrated non-diffuse repertoire organization at the whole-organ level. However, the whole-organ measurements integrate vascular, stromal, and parenchymal contributions into a single composite readout. Therefore, they can reveal the endpoint organ association but cannot determine which tissue compartments contribute to the observed repertoire structure. This limitation motivated direct comparison of capillary- and parenchyma-associated selection profiles.

Compartment resolution did more than add anatomical detail; it changed the information obtainable from systemic *in vivo* phage selection. The capillary-associated repertoire showed a comparatively continuous and overlapping organization, whereas the parenchyma-associated repertoire contained a compact shared region together with organ-associated extensions and peripheral structures. UMAP and HDBSCAN were used here as exploratory tools to visualize and partition peptides according to similarity in their sample-wise enrichment profiles, not to infer cell identities or direct molecular binding states. In this sense, the analytical logic was adapted from single-cell transcriptomics, but the measured object remained the repertoire of peptides recovered after *in vivo* selection. Dissociation-based single-cell workflows provide direct information about cellular composition and state, but enzymatic processing can alter cell-surface marker recovery, cellular composition, and transcriptional state (14–16). By design, dissociated-cell assays also do not preserve the intact anatomical context in which systemic peptide exposure and recovery occur. Conversely, LCM enabled morphology-guided sampling of anatomically defined regions without enzymatic tissue dissociation. Nevertheless, LCM-defined fractions do not establish molecularly pure or single-cell populations, and adjacent structures may contribute to the recovered signal. Therefore, we interpret these data as capillary- and parenchyma-associated enrichment profiles rather than cell-type-specific binding maps. Within these boundaries, the differences between bulk and compartment-resolved datasets indicate that resolving vascular and parenchymal compartments provides qualitatively distinct repertoire information.

Motif-centered analyses provided sequence-level depth to the compartment-resolved landscapes. The representative STLHQ-defined neighborhood was sharply bounded yet retained limited sequence variation, whereas the STFHQ-defined neighborhood showed pronounced convergence toward a single detected sequence within the observed repertoire. This contrast was supported by complementary PSSM, bootstrap, tiered-perturbation, and entropy analyses and therefore was not based on sequence-logo visualization alone. These two representative neighborhoods illustrate a range of local sequence-tolerance regimes, from bounded variation to near single-sequence convergence, but they should not be interpreted as defining the behavior of the entire recovered repertoire. Importantly, the compartment in which a medoid was identified does not imply exclusive compartmental assignment of its corresponding motif-defined set. A medoid represents a geometry-derived cluster center within a specific embedding, whereas enrichment of the STLHQ- and STFHQ-defined sets extended across both capillary- and parenchyma-associated samples. Thus, these neighborhoods were compartment-anchored but not compartment-exclusive. The two motif-defined sets also showed concordant enrichment at a restricted subset of anatomical sample labels. This convergence may identify tissue contexts in which capillary- and parenchyma-associated selection signals overlap, although it does not establish a distinct biological category or identify the molecular basis of the shared enrichment.

The divergence between bulk and compartment-resolved analyses was informative rather than contradictory. In the bulk cerebrum dataset, multi-organ digital subtraction identified a cerebrum-restricted upper-ranking regime by comparing enrichment in the cerebrum with the strongest signal observed across the remaining organ panel. This analysis quantified endpoint organ selectivity but could not determine whether the underlying signals originated from vascular or parenchymal compartments. Conversely, peptides highlighted by compartment-resolved analysis were not necessarily prioritized in the bulk-organ ranking because whole-organ profiles combine contributions from multiple tissue environments. EVGTARY provided an exploratory example of this distinction. It was not among the top-ranked peptides in the bulk cerebrum analysis but was enriched in cortex-associated parenchymal samples and localized near the medoid of a brain-associated local cluster. Following incorporation into a BPEI-based polyplex, EVGTARY was associated with increased DAPI-positive area-normalized cortical DsRed fluorescence compared with BPEI polyplex alone in the mouse-level ROI analysis. This observation supports the use of compartment-resolved landscapes for nominating experimentally testable candidates that may be deprioritized by bulk-organ ranking, though it does not establish sequence-specific cortical targeting.

Several limitations define the interpretation of these findings. First, in vivo phage display integrates systemic exposure, tissue recovery, bacterial amplification and propagation, and sequencing; the resulting profiles cannot be attributed solely to molecular binding within tissues. Same-lot naïve-library profiling did not detect the exact STLHQEL, STFHQKL, or EVGTARY sequences or the examined STLHQxx and STFHQxx motif matches in either profile from the same unamplified input-library lot, weakening the possibility that these features reflected high detectable input abundance. However, the naïve profiles were not used for selected-to-input normalization or landscape recalculation, and broader library or propagation effects cannot be excluded. Second, the complete body-wide organ–compartment selection series was not replicated in an independent selection series. The present dataset should therefore be interpreted as a discovery landscape, with the reproducibility of both repertoire-level patterns and individual candidate peptides remaining to be established through targeted independent experiments. Aggregated sequencing-read counts also do not estimate between-animal biological variability, and the molecular receptors and mechanisms underlying the observed organization remain unidentified. Finally, the EVGTARY experiment was exploratory, included four mice per group, lacked a sequence-scrambled peptide control, and relied on ROI-based fluorescence rather than an independent molecular assay. Sequence-independent effects of peptide conjugation, carrier context, formulation, or physicochemical properties therefore remain possible, and the affected step in the delivery cascade is unknown. Future work will require targeted replication, candidate-specific controls, quantitative assays, and mechanistic studies.

## 4. Conclusion

This study provides a systematic repertoire-level view of peptide recovery after systemic *in vivo* phage selection across whole-organ, capillary, and parenchymal scales. The recovered repertoires showed structured, compartment-dependent organization, while representative motif-defined neighborhoods exhibited local sequence-tolerance regimes ranging from bounded variation to near single-sequence convergence. Compartment resolution also yielded information that was qualitatively different from whole-organ endpoint ranking and enabled nomination of an experimentally testable candidate that was not prioritized in bulk analysis. Together, these findings shift the analytical unit of *in vivo* phage display from isolated organ-associated hits toward the organization of recoverable peptide repertoires and support a model of tissue-constrained peptide recovery. The resulting normal-tissue landscape provides a reference point for future mechanistic studies and candidate-specific evaluation without presupposing direct receptor binding, targeting specificity, or clinical utility.

## 5. Experimental Section

### *In vivo* biopanning and organ collection

#### Mouse model

##### Animals and housing

C57BL/6J msSlc mice of both sexes, aged 7–12 weeks, were supplied by Shimizu Laboratory Supplies Co., Ltd. (breeder: Japan SLC, Inc.). The animals were maintained under standard housing conditions with a 12-h light/12-h dark cycle and were given free access to standard laboratory chow and water.

The number of mice used varied depending on the stage of *in vivo* phage display biopanning. For the first round, approximately three mice were allocated per target organ to ensure sufficient recovery. For the second and third rounds, one to two mice were typically used per organ. After the final available selection round, the enriched phage preparation was administered to one to two mice for terminal collection of organs. For all organs except urinary bladder, the final available round corresponded to round 3; for urinary bladder, the final available round corresponded to round 2.

##### Perfusion protocol

For all *in vivo* phage display experiments, 2 × 10^11^ pfu of phage library in 200 µL was administered via the tail vein and allowed to circulate for 5 min before tissue collection. For terminal tissue collection, mice were deeply anesthetized with medetomidine (0.3 mg kg⁻¹), midazolam (4.0 mg kg⁻¹), and butorphanol (5.0 mg kg⁻¹), administered intraperitoneally, followed by PBS perfusion and organ collection. Before organ collection, mice were perfused with sterile PBS (−) for 5 min to reduce residual blood-associated and circulating phage. For collection of circulating blood, blood was obtained by cardiac puncture before PBS perfusion.

##### Ethics statement

All animal procedures were approved by the Institutional Animal Care and Use Committee of Shiga University of Medical Science (approval numbers: 2018-4-11 (H1) and 2021-1-4 (H1)) and were conducted in accordance with the applicable institutional guidelines and regulations. All efforts were made to minimize animal suffering.

##### Phage library

A Ph.D.-C7C Peptide Library Kit (New England Biolabs, Ipswich, MA, USA) was used for the *in vivo* phage display experiments. Escherichia coli K-12 ER2738 (New England Biolabs, #E4104S) was used for phage propagation and titer determination.

On the day of phage administration, *Escherichia coli* ER2738 cells were cultured in LB broth at 37°C with shaking (150 rpm) until the culture reached an OD600 of approximately 0.5–0.6. The phage preparations used for mouse injection were diluted in TBS buffer to the desired administration concentration.

#### *In vivo* phage display biopanning

##### Bulk biopanning workflow

For bulk *in vivo* biopanning, phage administration, 5-min circulation, circulating blood collection, and PBS perfusion were performed as described in the Perfusion protocol section. After perfusion, target organs were harvested, washed with PBS (−), weighed, and homogenized in 500 µL of DMEM supplemented with protease inhibitor.

For titer determination, tissue homogenates were serially diluted in TBS. Aliquots of each dilution, including the undiluted sample, were added to *Escherichia coli* ER2738 suspensions and incubated for 5 min at room temperature to allow phage infection. The infected bacteria were mixed with top agar, plated on LB/IPTG/X-gal plates, and incubated overnight at 37°C. Blue plaques were counted the following day to estimate phage recovery. *In vivo* phage display biopanning was generally performed for three rounds, with amplification steps conducted between rounds as described below. Urinary bladder was an exception and was analyzed using the second-round enriched sample as the final available selected sample.

##### Phage recovery and amplification

The plates inoculated with undiluted samples were overlaid with SM buffer and incubated at 4°C overnight. The phage-containing SM buffer was then recovered and stored at 4°C until amplification.

For amplification, the recovered phage-containing SM buffer was added to *Escherichia coli* ER2738 cultures and incubated at room temperature for 5 min to allow phage infection. Infected cultures were expanded in LB broth supplemented with MgSO4 at 37°C with shaking for 1 hr.

After amplification, chloroform was added, and the cultures were centrifuged to remove bacterial debris. Phage particles in the supernatant were precipitated using PEG/NaCl at 4°C overnight, pelleted by centrifugation, and resuspended in TBS. Phage titers of the amplified preparations were determined using the plaque assay described above for titer determination. Amplified phage preparations were adjusted to the desired concentration and used for subsequent rounds of biopanning.

##### Sample collection for sequencing

After the final round of biopanning, the tissues were collected for downstream sequencing analysis. For bulk analysis, the tissue samples were directly processed for DNA extraction.

In parallel, selected tissues were embedded in OCT compound, frozen, and sectioned for laser capture microdissection (LCM) to enable compartment-resolved analysis. Tissues were allocated for bulk and LCM processing without assuming identical cellular compositions between the two preparations.

Laser capture microdissection (LCM) was performed by using an Arcturus XT Laser Capture Microdissection System (Thermo Fisher Scientific, USA). Frozen tissue sections (4–5 μm) were prepared by using a cryostat and mounted onto glass slides. Sections were stained with hematoxylin and eosin (H&E), and target cells were selected microscopically and captured using Arcturus CapSure Macro LCM Caps (Thermo Fisher Scientific, USA).

The number of captured cells varied depending on the sampled tissue region and was therefore not fixed across samples. DNA was extracted from the captured cells by using the QIAamp DNA Micro Kit (QIAGEN, Hilden, Germany) according to the manufacturer’s protocol.

Lists of parenchymal LCM samples are provided in Table S2.

DNA was extracted from bulk tissues using a QIAamp DNA Mini Kit (QIAGEN) and a list of bulk organs is provided in Table S1.

#### DNA extraction and sequencing

##### PCR amplification and library preparation

Phage-derived DNA from the bulk samples (SM buffer collected after panning) and DNA extracted from the LCM-captured samples were used as templates for PCR amplification. The primers were designed to include 5′ overhang adaptor sequences followed by phage-specific sequences. The primer sequences used were as follows: Forward: 5′-TCGTCGGCAGCGTCAGATGTGTATAAGAGACAGttggtgccggcaac-3′ Reverse: 5′-GTCTCGTGGGCTCGGAGATGTGTATAAGAGACAGgccctctatagttagc-3′ PCR reactions were performed by using the following reaction mixture: 5 μl of PCR buffer, 4 μl of dNTP mix, 1 μl each of forward and reverse primers (10 μM), 0.3 μl of Ex Taq polymerase (Takara Bio, Kusatsu, Japan) and template DNA (1 μl for SM buffer samples or 5 μl for LCM samples), with nuclease-free water added to a final volume of 50 μl.

The thermal cycling conditions were as follows: initial denaturation at 94°C for 3 min; 30–45 cycles of 94°C for 30 s, 60°C for 30 s, and 72°C for 30 s; and a final extension at 72°C for 7 min.

The PCR products were separated by electrophoresis on 1.5% agarose gels, and DNA fragments of the expected size were excised under blue LED illumination and purified using the NucleoSpin Gel and PCR Clean-up kit (Takara Bio).

##### Next generation sequencing

Purified DNA libraries were subjected to sequencing on an Illumina MiSeq platform (outsourced to Hokkaido System Science, Sapporo, Japan). The sequencing reads were processed to obtain the corresponding 7-mer peptide sequences.

#### NGS data preprocessing

##### Raw read processing

Raw FASTQ files were used for public sequencing-read deposition. For peptide-level preprocessing, service-derived nucleotide sequence tables in comma-separated value (CSV) format were used and translated into peptide sequences by using a custom in-house pipeline. Only reads that could be unambiguously translated into full-length 7-amino-acid peptides in the correct reading frame and without premature stop codons were retained. Truncated sequences and reads containing ambiguous or incomplete translations were discarded.

The retained reads were then collapsed at the peptide level to generate count tables in which each row corresponded to a unique 7-mer peptide and each column represented a sample-level count profile. These peptide-level count tables served as the input for all downstream analyses.

##### Filtering thresholds and exception handling

For the main analyses, extremely low-abundance peptides were removed before downstream processing. Sequences with fewer than 20 total reads after the final available enriched round of *in vivo* biopanning were excluded as low-abundance sequences with the rationale that specifically enriched phage clones would be expected to accumulate across iterative selection rounds. This threshold was applied consistently to the bulk, capillary, and parenchymal datasets before counts per million (CPM) normalization, UMAP embedding, and clustering analyses.

An exception was made for digital subtraction analysis, in which all observed 7-mer sequences were retained regardless of abundance to capture potentially informative early-stage enrichment signals that may not yet have exceeded the filtering threshold after the final available enriched round.

##### Normalization and data transformation

For count-based comparisons, raw peptide counts were normalized to the CPM, defined as follows:

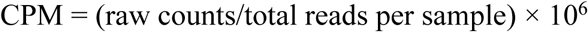

The CPM values were used for rank-curve analysis, digital subtraction, sequence enrichment evaluation, and motif abundance comparisons.

For manifold learning and other distributional analyses, raw peptide count matrices were transformed by using log1p. The values were then normalized across each peptide to emphasize the relative organ- or sample-wise distribution patterns independent of the absolute read depth, and features with zero variance across samples were removed before downstream embedding.

For distributional comparisons between the capillary and parenchymal layers, empirical cumulative distribution functions (ECDFs) were computed separately for each layer from log1p-transformed CPM values across all peptide–sample observations. For a given value x, the ECDF was defined as the fraction of observations with log1p(CPM) ≤ x.

##### Sequence aggregation and deduplication

All downstream analyses were performed at the level of unique peptide sequences by using peptide-level count tables with aggregated counts for each sample. No additional read-level deduplication or PCR-duplicate removal was applied after this step.

##### Organ mapping

Each sequencing dataset (bulk, parenchyma, and capillary) was annotated by using a metadata table specifying the organ identity of each sample. The parenchymal and capillary datasets were derived from laser capture microdissection and therefore included multiple spatial subregions from the same organ. These subregions were retained as separate entries rather than merged, in order to preserve the spatial resolution and within-organ heterogeneity.

The compartment labels (parenchyma or capillary) were retained to distinguish the tissue compartments, whereas the bulk samples consisted of a single sample per organ and therefore did not include compartment labels. Organ names and physiological system annotations were harmonized across the bulk, capillary, and parenchymal datasets to enable cross-dataset comparisons.

For the capillary dataset, pancreas-related labels 11 and 37 were evaluated during preprocessing because they represented overlapping pancreas-associated sequence groups. Based on this comparison, label 37 was collapsed into label 11 for downstream organ assignment, and the resulting curated mapping was used for subsequent principal component analysis (PCA)/UMAP analysis.

#### Same-lot naïve library background profiling

##### Naïve-library datasets and provenance

Raw FASTQ data were available for two naïve-library sequencing datasets, L35742 and L35743. Both datasets were generated from the same naïve phage library lot used as the input library for the mouse *in vivo* selection experiments. No intervening *E. coli* amplification of the phage library was performed before sequencing. Thus, L35742 and L35743 represent two sequencing/read-mode profiles of the same unamplified naïve library rather than independent library lots, independent biological replicates, or naïve-versus-amplified experimental conditions. The datasets were analyzed separately to characterize the detectable sequence background of the same-lot input library.

##### Processing and enumeration of naïve-library sequences

The merged peptide-count table for L35742 and L35743 was processed independently of the *in vivo*–selected datasets. Peptide sequences were converted to uppercase, and only full-length 7-mer sequences composed exclusively of the 20 standard amino acids (ACDEFGHIKLMNPQRSTVWY) were retained. Entries of other lengths and sequences containing nonstandard or ambiguous characters were excluded. No additional read-count threshold was applied after loading the service-derived peptide-count table. All valid 7-mer sequences represented in the supplied profiles were retained for exact-sequence and motif queries.

Identical valid 7-mer sequences were represented as individual peptide-level rows with separate count values for L35742 and L35743. CPM values were recalculated independently for each profile by using the total number of valid 7-mer reads in that profile as the denominator. Peptides were ranked separately within L35742 and L35743 in descending order of raw read count. For conservative characterization of the detectable input-library background, the maximum CPM observed across the two profiles was additionally calculated for each peptide. L35742 and L35743 were not pooled for CPM or rank calculation. Of 19,816 peptide-table entries, 19,653 satisfied the valid 7-mer criteria. The valid 7-mer read totals used as the CPM denominators were 80,588 for L35742 and 76,158 for L35743.

##### Exact-sequence and motif queries

The naïve-library profiles were queried for the exact 7-mer peptide sequences STLHQEL, STFHQKL, and EVGTARY. Exact-sequence detection was defined as complete amino-acid identity across all seven positions. Motif sequence searches were performed for all valid 7-mer sequences matching the anchored patterns ^STLHQ..$ and ^STFHQ..$, corresponding to peptides beginning with STLHQ or STFHQ followed by any two standard amino acids.

Queries were performed separately for L35742 and L35743 by using the unthresholded valid 7-mer count tables. An exact peptide or motif member was classified as detected when its raw read count was greater than zero in the corresponding profile. A feature was classified as not detected only when no matching valid 7-mer read was present in that profile. Raw read counts and profile-specific CPM values were calculated for descriptive assessment.

##### Relationship to downstream landscape analyses

The same-lot naïve-library profiles were used only for descriptive input-background assessment. They were not incorporated into the normalization of the *in vivo*–selected datasets and were not used to filter peptide sequences before downstream analysis. The UMAP embeddings, HDBSCAN clustering, digital subtraction, motif analyses, medoid calculations, and candidate-ranking procedures were therefore not recalculated or modified on the basis of the naïve-library profiles.

No selected-to-naïve enrichment ratios or naïve-corrected abundance measures were calculated. Accordingly, this analysis was interpreted as a targeted assessment of whether the major selected peptide features were detectable as prominent components of the sequenced same-lot input-library background, rather than as direct naïve normalization of the selected peptide landscapes.

#### Digital subtraction analysis

##### Data normalization

For digital subtraction analysis in the bulk dataset, all observed 7-mer peptides from the first and final available enriched selection rounds were retained without abundance-based filtering. For all organs except urinary bladder, the final available enriched round corresponded to round 3; for urinary bladder, the final available enriched round corresponded to round 2. The raw counts for each organ were normalized to CPM to enable comparison across samples with different sequencing depths.

##### Definition of the enrichment and subtraction score

Organ-specific enrichment scores were calculated from the CPM-normalized counts as log_2_ fold changes between the final available enriched round and the first selection round:

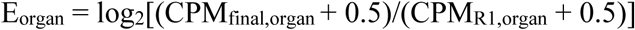

where CPMfinal denotes CPM at round 3 for all organs except urinary bladder and CPM at round 2 for urinary bladder.

A pseudocount of 0.5 was added to avoid undefined values for zero counts and to stabilize the fold-change estimates for low-abundance peptides. To restrict the analysis to positive enrichment signals, nonpositive enrichment values (log_2_ FC ≤ 0) were set to zero before calculation of the subtraction score.

For the cerebrum-specific analysis, a digital subtraction score was defined as follows: subtraction score = E_cerebrum_ – max (E_other organs_) where E_other organs_ denotes the maximum enrichment score observed across all other mapped organs.

##### Rank curve analysis

Peptides were ranked by subtraction score to examine the distribution of cerebrum-specific enrichment across the bulk dataset. Rank-curve analysis was used to assess the extent of cerebrum-exclusive enrichment and its transition to peptide sequences showing shared enrichment across multiple organs.

##### Visualization of organ specific enrichment profiles

Heatmaps were generated by using the organ-specific enrichment values (E_organ_) for the peptides ranked within the top 50 according to the cerebrum subtraction score for visualization. For each peptide, the enrichment values in the cerebrum, liver, and skeletal muscle were extracted from the bulk sequencing data. The liver and skeletal muscle were included as representative nonneural organs to assess whether strong enrichment was restricted to the cerebrum rather than accompanied by comparable enrichment in off-target organs.

##### Rationale for digital subtraction

Digital subtraction was designed to formalize organ selectivity by explicitly contrasting enrichment in the target organ with the maximum enrichment observed across all other organs within the same bulk dataset. This approach allows target-organ enrichment and potential off-target enrichment to be evaluated within the same scoring scheme.

In this study, digital subtraction was applied to the bulk dataset using the cerebrum as the target organ to quantify organ selectivity from whole-organ sequencing data. Because bulk samples inherently contain mixed vascular and parenchymal signals, vascular and parenchymal contributions were evaluated separately using laser-capture microdissection-derived datasets, as described in the compartment-resolved analyses.

##### Embedding and motif discovery

Embedding and clustering analyses were informed by high-dimensional analytical workflows widely used in single-cell transcriptomics, but were adapted here to organ-and compartment-resolved peptide enrichment profiles rather than cellular transcriptomes. Accordingly, UMAP and HDBSCAN were used as exploratory tools to visualize and partition peptide sequences according to similarity in enrichment profiles, rather than to infer cell types or transcriptional states.

##### UMAP parameters

UMAP embeddings were generated in an unsupervised manner from the top 10 principal components derived from the log1p-transformed, row-normalized peptide count matrix, without the use of organ or compartment labels (*17*). Unless otherwise noted, UMAP was run with the following parameters: n_components = 2, n_neighbors = 30, min_dist = 0.2, metric = ‘cosine’, and random_state = 42.

##### Embedding space definition

All dimensionality reduction steps (PCA and UMAP) were performed in a fully unsupervised manner; the system and organ labels were not used during embedding construction and were applied only for visualization and post hoc interpretation.

Pairwise distances in UMAP space were interpreted as approximate reflections of dissimilarity among the organ-wise enrichment profiles of the peptides. Local neighborhoods were interpreted as preservation of the proximity relationships present in the PCA-reduced space derived from the normalized count matrix. Thus, regions of the embedding do not correspond to meaningful positional axes but instead represent peptides with similar organ-resolved enrichment patterns.

##### Robustness to PCA dimensionality

To assess whether the embedding and downstream clustering were sensitive to PCA dimensionality, the analysis was repeated by using 10, 20, and 41 principal components, while keeping all of the UMAP and HDBSCAN parameters fixed. The upper value of 41 corresponded to the full dimensionality of the organ feature space in the bulk dataset.

The concordance between cluster assignments obtained under different PCA dimensionalities was quantified by using the adjusted Rand index (ARI) and normalized mutual information (NMI). These analyses indicated that the overall clustering structure was stable across PCA dimensionalities, supporting the robustness of the selected PCA setting.

##### Embedding structure assessment

The intrinsic dimensionality was estimated before the UMAP projection to assess the effective dimensionality of the normalized peptide count space (*18*). Estimates were obtained by using the Levina–Bickel maximum-likelihood estimator with k = 10 nearest neighbors; self-neighbors were excluded from the distance calculations. For datasets containing more than 5,000 peptides, a random subset of 5,000 peptides was used for computational efficiency (random_state = 0). This procedure provided a quantitative assessment of the dimensional structure prior to UMAP embedding.

To assess the robustness to stochastic initialization, the UMAP embeddings were recomputed by using different random seeds (random_state = 0, 42, and 123) while keeping all of the preprocessing steps and UMAP parameters identical.

To assess whether the observed embedding structure could be explained by chance, two null controls were constructed. In the label-permutation control (null model A), the embedding coordinates were kept fixed, and the system labels were randomly permuted for visualization to assess the apparent label coherence under random assignment. In the feature-wise shuffle control (null model B), each feature column in the peptide-by-feature matrix was independently permuted prior to PCA and UMAP, thereby disrupting peptide–feature correspondence while preserving marginal feature distributions; the same PCA and UMAP pipeline was then applied.

To quantify the shared-core structure in the bulk UMAP embedding, the embedding center was defined as the centroid of the real (non-shuffled) coordinates, and the Euclidean distance from each point to this center was calculated. The core radius was defined as the 10th percentile of these distances in the real embedding. The shared-core density was quantified as the fraction of points falling within this radius (core-in fraction), and the global spread was quantified as the median distance from the center. For direct comparison, the same center and radius were applied to the feature-wise shuffled null embedding (null model B). Robustness to the choice of core radius was assessed by repeating the analysis with thresholds defined by the 5th and 15th percentiles of the distances in the real embedding.

##### HDBSCAN clustering and parameter optimization

HDBSCAN clustering was performed on the UMAP embedding to identify density-defined peptide groups (*19*). The clustering separability was assessed by using the silhouette score, computed on the two-dimensional UMAP embedding with Euclidean distance. This metric was used to characterize differences in the cluster compactness and overlap across datasets rather than as an optimization objective.

The minimum cluster size parameter was selected through a parameter sensitivity analysis that evaluated the clustering behavior and organ-level consistency across a range of values. The clustering results obtained with different minimum cluster sizes were compared against the organ labels by using the adjusted Rand index (ARI) and normalized mutual information (NMI). Based on this analysis, min_cluster_size = 50 was selected as a parameter setting that maintained organ-level consistency without excessive cluster fragmentation.

The final clustering model used HDBSCAN with min_cluster_size = 50, metric = ‘euclidean’, and cluster_selection_method = ‘eom’; these parameters were applied to the two-dimensional UMAP embedding generated by using a cosine distance metric.

A parameter sensitivity analysis was performed by using the bulk dataset, which contained the most heterogeneous organ composition. The same parameter set was subsequently applied to the parenchymal and capillary datasets without further tuning.

##### Cross-compartment overlap analysis and medoid extraction

To identify peptide clusters shared across datasets and tissue compartments, cross-compartment relationships were assessed between the clusters derived from the capillary and parenchymal datasets using the same UMAP and HDBSCAN parameter settings. The cluster-level peptide overlap was evaluated by using a one-sided hypergeometric test. For a given cluster pair A and B, the universe (|U|) was defined as the union of unique peptide sequences observed in all non-noise clusters from the two datasets under comparison. The statistical significance of overlap was computed by using the survival function of the hypergeometric distribution.

Cluster pairs exhibiting high overlap enrichment (observed-to-expected ratio > 5) together with nominal statistical significance (P < 0.01) were retained as candidate shared cluster pairs in an initial screening step for downstream analyses.

For each cluster within the retained cross-compartment pairs, a representative peptide sequence was identified by medoid extraction in the two-dimensional UMAP embedding space. The pairwise Euclidean distances were computed among the peptide coordinates (UMAP1, UMAP2) within each cluster, and the medoid was defined as the peptide that minimizes the sum of distances to all other peptides in the same cluster. The resulting medoid sequences were carried forward for motif solution-space definition and downstream analyses.

#### Compartment resolved motif analysis

##### Motif definition and sequence logo analysis

Peptide sequences recovered from *in vivo* selection were stratified by tissue compartment (capillary or parenchyma) and aggregated across anatomical sample labels within each compartment for motif analysis. Sequence logo analyses were performed in a read-weighted manner unless otherwise specified.

The resulting medoid sequences, STLHQEL and STFHQKL, were used as anchors for motif definition. Peptides identical to each medoid sequence were first identified via direct string comparison. Subsequently, relaxed motif patterns were defined by using regular expression-based matching, allowing variation at the C-terminal two positions while preserving the N-terminal core. These baseline motif definitions, designated Tier 0, were specified by the patterns ^STLHQ..$ and ^STFHQ..$. Pattern matching was implemented in Python using regular expressions and applied to both the capillary and parenchymal peptide datasets. All of the peptides matching each pattern were collected and used as inputs for subsequent analyses.

For each Tier 0 motif-defined peptide set within each compartment, position-specific amino acid counts were computed in a read-weighted manner after the reads for identical peptide sequences were aggregated. Specifically, the read counts were first summed for each unique 7-mer, and a position-by-amino-acid count matrix was then constructed from the resulting sequence-level counts. For the sequence logos shown in Fig. 3b, the background amino acid frequencies were estimated separately for each tissue compartment by using all 7-mer peptides observed in that layer, weighted by read count. A pseudocount of 0.5 was added to both the background and position-specific counts. The position-specific scoring matrices (PSSMs) were calculated as log_2_-odds values by comparing the read-weighted amino acid frequency at each position with the corresponding read-weighted background frequency of the same layer. Sequence logos were generated from these background-corrected PSSMs by using the Logomaker Python package (*20*). For visualization in Fig. 3b, only the positive log_2_-odds values were displayed, with negative values set to zero.

##### Motif abundance heatmap

Peptides were identified by using the motif definitions described above and assigned to STLHQ- or STFHQ-matched peptide sets. For each motif-defined set, the CPM values were aggregated according to the anatomical sample labels and tissue layer (capillary or parenchyma) by summing the CPM across all matched peptides. The anatomical sample labels retained the original LCM naming convention, typically consisting of an organ identifier together with a more specific subregional or cell-associated label. The aggregated values were then transformed according to log1p (CPM).

Heatmaps were generated from the sample-by-layer matrices where rows corresponded to anatomical sample labels and columns to tissue layers. Missing capillary or parenchymal entries for a given sample label were filled with zeros. For each motif-defined set, sample labels were ranked by the maximum log1p (CPM) observed across the two layers, and the top 15 entries were displayed. For the top-ranked heatmaps shown in Fig. 3C, color scales were independently normalized within each motif-defined set and clipped at the 95th percentile of nonzero values. For the all-organ heatmaps shown in Fig. S12, the corresponding color scales were clipped at the 99th percentile of nonzero values.

##### Position specific scoring matrix and PSSM heatmap

For the PSSM heatmaps shown in Fig. 3d, position-specific scoring matrices were constructed for the STLHQ and STFHQ motif peptide subsets. Read-weighted position-specific amino acid counts were computed from the motif subset table, and amino acid frequencies were calculated after identical peptide sequences were represented by their read counts. Log₂-odds scores were calculated relative to a uniform 20-amino-acid background, with a background frequency of 1/20 assigned to each amino acid. No pseudocount was added for this PSSM heatmap calculation, and non-finite log₂-odds values arising from zero observed frequencies were set to zero before visualization. Unlike the compartment-stratified sequence logos shown in Fig. 3b, the PSSM heatmaps were generated from the full matrices, including both positive and negative log₂-odds values, and visualized using a zero-centered diverging color scale to show position-specific amino acid enrichment and depletion relative to the uniform background.

##### Anatomical-site-resolved enrichment analysis

Anatomical-site-resolved enrichment of the STLHQ- and STFHQ-defined peptide sets was evaluated using laser-capture microdissection-derived capillary and parenchymal peptide datasets. For each motif-defined peptide set, motif-associated read counts were aggregated according to the anatomical sample labels assigned to each LCM-captured region. For each anatomical sample label, an enrichment ratio was calculated as the relative abundance of the motif-associated reads within the anatomical sample label divided by the global relative abundance of the same motif-associated reads across the combined LCM organ panel.

Statistical significance was assessed using a one-sided Fisher’s exact test, with the alternative hypothesis set to greater. For each anatomical sample label, a 2 × 2 contingency table compared motif-associated and non-motif reads within that anatomical sample label with the corresponding reads pooled across all other anatomical sample labels in the same layer. The P values were corrected for multiple testing across anatomical sample labels by using the Benjamini–Hochberg false discovery rate (FDR) procedure (*21*). Enrichment ratios and adjusted q values were used for downstream interpretation and visualization. For plots showing −log10(q), the q values returned as zero due to numerical underflow were clipped to 1×10⁻^300^.

##### Concordant motif enrichment across anatomical sample labels

To identify anatomical sample labels exhibiting concordant enrichment of both motif-defined sets, anatomical sample labels showing significant enrichment in both the STLHQ-defined set and the STFHQ-defined set were first identified by using the significance criteria described in the previous section (q < 0.05). For each such anatomical sample label passing this criterion, the enrichment ratios for the two motif-defined sets were extracted. A combined concordance score, referred to as the motif convergence score, was calculated by multiplying the enrichment ratios of the two motif-defined sets within the same anatomical sample label. Anatomical sample labels were then ranked according to this score for visualization and comparison. The motif convergence score was used as a descriptive ranking metric and not as an independent statistical test.

##### Bootstrap analysis of the motif log odds

To assess the computational stability of motif-level positional constraints, nonparametric bootstrap resampling was performed at the sequence level. For each motif-defined peptide set and tissue layer (capillary or parenchyma), unique peptide sequences matching the predefined motif pattern were resampled with replacement for 2,000 bootstrap iterations (random seed = 123). Sampling probabilities were proportional to raw read counts to preserve the original abundance structure.

For each bootstrap sample, the position-specific log_2_-odds scores were recalculated using a pseudocount of 0.5 relative to a background distribution defined by all filtered 7-mer peptides in the corresponding layer. For each motif position, the mean log_2_-odds score and 95% bootstrap interval, defined by the 2.5th and 97.5th percentiles of bootstrap replicates, were calculated.

These statistics were computed for all seven positions; however, interpretation of the bootstrap-derived variability focused on positions 6 and 7 because positions 1–5 were invariant by construction under the Tier 0 motif definitions and therefore yielded negligible bootstrap variability.

##### Tiered perturbation analysis

For each motif-defined set, a read-dominant reference sequence was operationally defined as the peptide sequence with the highest read abundance: STLHQKL for the STLHQ-defined set and STFHQKL for the STFHQ-defined set. Tiered perturbation analysis was then performed on sequences pooled across anatomical sample labels within each tissue compartment (capillary or parenchyma) by using the read-weighted counts as described above.

To systematically probe the sequence tolerance around each reference motif, the additional perturbation tiers were defined using predefined regular expression patterns. Amino acid substitutions allowed in higher tiers were selected to represent conservative or motif-neighborhood substitutions rather than arbitrary sequence changes.

For the STLHQ-defined set, tiers were defined as follows:

Tier 1: ^ST[IVL]HQ..$

Tier 2: ^ST[TVLI]HQ..$

Tier 3: ^[ST][TL][IVL]HQ..$

Tier 4: ^STL[HNY]Q..$

For the STFHQ-defined set, only Tier 1 was defined (^ST[FWY]HQ..$) because candidate higher-tier patterns did not yield additional matched variants and were therefore not analyzed further.

Because subtle tier-dependent differences at positions 6 and 7 were not readily compared from sequence logos alone, read-weighted position-specific log₂-odds values at these positions were summarized as bar plots to enable direct quantitative comparison across tiers and tissue compartments.

##### Shannon entropy analysis

To quantify sequence-level diversity under tiered perturbation of the STLHQ motif family, read-weighted Shannon entropy was calculated for each predefined STLHQ tier (Tier 0, Tier 3 and Tier 4) separately in the capillary and parenchymal datasets. Peptides matching each tier-specific regular expression were extracted from the sequence-level read-count table. For each tier and tissue layer, the read fraction of peptide i was calculated as pi = ri / Σri, where ri denotes the read count of peptide i within the corresponding tier and layer. Shannon entropy was then calculated as H = −Σi pi log_2_ pi using peptides with pi > 0. No pseudocount was added for entropy calculation. Tiers with no matched peptides were assigned NA. Entropy values were visualized as bar plots to compare read-weighted sequence diversity across tiers and tissue layers.

#### Candidate selection and *in vivo* testing

##### Peptide selection criteria

To select a candidate peptide for *in vivo* testing, we used a multistep computational prioritization strategy based on embedding geometry and anatomical specificity. First, peptides enriched in cortex-associated LCM samples were identified. Candidates were then prioritized if they occupied peripheral positions in the unified embedding of the capillary and parenchymal datasets, distinct from the dominant STLHQ- and STFHQ-associated motif regions.

Next, local cluster structure was assessed by using HDBSCAN, and candidates were further prioritized if they localized near the medoid of their assigned cluster, indicating that they represented cluster-central representatives rather than isolated outliers. Finally, candidates exhibiting parenchymal enrichment together with low signal across nontarget organs were prioritized for exploratory *in vivo* assessment. On the basis of these criteria, EVGTARY was selected for downstream *in vivo* experiments.

##### EVGTARY-BPEI polyplex administration and cortical signal assessment

Branched polyethyleneimine (BPEI; nominal Mw 25,000, Mn approximately 10,000; Sigma- Aldrich, Tokyo, Japan) was purchased and used as received. The dsRed plasmid (AVS-siRNA-dsRed) was kindly provided by Dr. Oka (Baylor College of Medicine, TX, USA). A construct containing the CMV promoter and dsRed coding sequence was used in this study.

An azide-functionalized synthetic ligand containing the selected EVGTARY core flanked by cysteine residues (Cys–EVGTARY–Cys) was conjugated to alkyne-modified BPEI by copper(I)-catalyzed azide–alkyne cycloaddition. The alkyne-modified BPEI was prepared by derivatization with 5-hexynoic acid, and conjugation to the azide-functionalized peptide was performed using CuI and N,N-diisopropylethylamine in dimethyl sulfoxide. Formation of the alkyne-modified BPEI and the peptide conjugate was assessed by ^1^H NMR spectroscopy, with estimated incorporation levels of approximately 4–5% and 2%, respectively. As a control polymer, benzyl-triazole–modified BPEI lacking peptide was prepared in parallel. The resulting polymers were mixed with dsRed plasmid DNA to form polyplexes through charge-mediated assembly. For gel retardation assays, the gel was stained with ethidium bromide and imaged under UV illumination using a UVP BioDoc-It/GDS-7900 system. All lanes shown were derived from a single unspliced gel, and a molecular-size marker was included. The amount of dsRed plasmid DNA was fixed at 200 µg for all formulation conditions and mixed with BPEI or EVGTARY–BPEI at a series of polymer-to-DNA equivalent ratios, expressed relative to plasmid DNA (= 1), and calculated on the basis of charge neutralization. Mixtures were prepared in water to a final volume of 200 µL and incubated for 30 min at room temperature before agarose gel electrophoresis. Based on these experiments, equivalent ratios of 1:50 for BPEI and 1:200 for EVGTARY–BPEI were selected for the subsequent *in vivo* studies.

Polyplexes corresponding to 2.04 µg of dsRed plasmid DNA were administered intravenously via the tail vein in a total volume of 200 µL per mouse under the selected formulation conditions. Mice were randomly allocated to the three treatment groups and sacrificed 4 days after injection and were perfusion-fixed with 4% paraformaldehyde. Brain tissues were harvested, cryosectioned at 20 µm thickness, and analyzed for dsRed signals in the cerebral cortex.

For quantitative image analysis, cortical sections were imaged at 400× magnification by a confocal laser microscope under identical acquisition settings (Leica TCS SP8X; Leica Microsystems GmbH, Wetzlar, Germany). Imaging and ROI quantification were performed with group identity blinded. ROI-based quantification of DsRed fluorescence was performed using ImageJ/Fiji on matched DsRed and DAPI channel images. Fixed-size regions of interest were placed within the cerebral cortex and applied to the corresponding DsRed and DAPI images. In the DsRed channel, integrated density was measured within each ROI. In the DAPI channel, images were converted to 8-bit grayscale and thresholded to determine the DAPI-positive area within the same ROI. The DAPI-positive area was calculated as the ROI area × (%Area/100). No mice, sections, microscopic fields, or data points were excluded, and no prespecified exclusion criteria were used.

DAPI-positive area-normalized DsRed fluorescence was calculated as DsRed integrated density divided by the DAPI-positive area. For each mouse, two microscopic fields were quantified and averaged before statistical analysis. The statistical unit was the mouse, with n = 4 mice per group. Normalized DsRed fluorescence was compared among groups using the Kruskal–Wallis test followed by Dunn’s post hoc test with Benjamini–Hochberg correction for multiple comparisons. Adjusted P values < 0.05 were considered statistically significant.

##### Computational implementation

All of the computational analyses were performed in Python using custom scripts. UMAP embeddings were generated with umap-learn (v0.5.9), and clustering analyses were performed by using HDBSCAN (v0.8.40). Additional statistical and numerical analyses were conducted with scikit-learn (v1.6.1), NumPy, SciPy, and pandas. Data visualization was performed by using matplotlib, and the sequence logos were generated with Logomaker.

##### AI-assisted computational support

ChatGPT (OpenAI) was used to assist with computational clarification, code drafting, and troubleshooting for analyses specified by the authors. The analysis objectives, parameters, and interpretation were determined by the authors. All AI-assisted code was reviewed, edited as necessary, executed, and verified by the authors, who take full responsibility for the computational analyses.

#### Statistical analysis

Statistical inference was limited to the cluster-overlap screening, anatomical sample label-resolved motif-enrichment analyses, and exploratory comparison of cortical fluorescence among treatment groups. Other quantitative analyses, including intrinsic-dimensionality estimation, adjusted Rand index, normalized mutual information, silhouette scores, empirical cumulative distribution functions, digital-subtraction scores, Shannon entropy, embedding-seed comparisons, null-model controls, and same-lot naïve-library background profiling, were used as descriptive or robustness assessments and were not interpreted as formal hypothesis tests unless otherwise specified.

Overlap between peptide clusters was evaluated using a one-sided hypergeometric test. For each pair of datasets, the peptide universe was defined as the union of unique peptide sequences assigned to all non-noise clusters in the two datasets. For each cluster pair with at least one shared peptide, the expected overlap was calculated as the product of the two cluster sizes divided by the peptide-universe size, and the observed-to-expected overlap ratio was calculated as an enrichment measure. Candidate cross-compartment cluster pairs were identified using an observed-to-expected ratio greater than 5 together with a nominal *P* value below 0.01. These nominal *P* values were used for exploratory screening and were not adjusted for multiple comparisons.

Anatomical sample label-resolved enrichment of the STLHQ- and STFHQ-defined peptide sets was assessed using one-sided Fisher’s exact tests with the alternative hypothesis set to greater. For each anatomical sample label, motif-matching and non-matching read counts were compared with the corresponding counts pooled across all other anatomical sample labels in the same tissue layer. *P* values were adjusted across LCM-derived anatomical sites within each motif-by-layer analysis using the Benjamini–Hochberg false-discovery-rate procedure, and adjusted *q* values below 0.05 were considered statistically significant. The motif convergence score was calculated as the product of the corresponding enrichment ratios among anatomical sample labels meeting the significance criterion in both component analyses. This score was used as a descriptive ranking metric and was not subjected to an additional statistical test. Because these analyses were based on aggregated sequencing-read counts, they quantified enrichment within the observed sequencing profiles rather than between-animal variability.

Computational stability of motif positional constraints was evaluated using 2,000 nonparametric bootstrap iterations. Within each motif-defined peptide set and tissue layer, unique peptide sequences were resampled with replacement using sampling probabilities proportional to their raw read counts. Position-specific log_2_-odds scores were recalculated for each bootstrap sample, and 95% percentile bootstrap intervals were defined by the 2.5th and 97.5th percentiles of the resulting distributions. A random seed of 123 was used. These intervals assessed stability within the observed peptide repertoire and were not interpreted as estimates of biological variability among animals.

For ROI-based quantification of cortical DsRed fluorescence, two microscopic fields were averaged within each mouse before statistical analysis, and the mouse was used as the statistical unit (*n* = 4 mice per group). Overall differences among the three treatment groups were evaluated using the Kruskal–Wallis test implemented with scipy.stats.kruskal. Pairwise Dunn comparisons were calculated from pooled ranks using scipy.stats.rankdata, with two-sided *P* values obtained from the standard normal distribution using scipy.stats.norm. The standard tie correction based on Σ(*t*³ − *t*), where *t* denotes the number of tied observations within each tie group, was included. All 12 mouse-level values were distinct, and the tie-correction factor was therefore 1.0. The three pairwise comparisons—plasmid only versus BPEI polyplex, plasmid only versus EVGTARY polyplex, and BPEI polyplex versus EVGTARY polyplex—were jointly adjusted using the Benjamini–Hochberg procedure. Adjusted *P* values below 0.05 were considered statistically significant. This proof-of-concept analysis was interpreted as exploratory. The ROI analysis was performed in Python 3.9.6 using NumPy 2.0.2, pandas 2.3.2, and SciPy 1.13.1.

## Acknowledgments

We thank Mr. Fumihisa Kojima for developing a program to convert raw sequencing output into peptide sequences, and Ms. Minako Matsumoto for synthesizing the EVGTARY–BPEI polyplex under the supervision of Y.F. We thank Ms. Yuki Nakae and Ms. Kyoko Furuhashi for their technical assistance with this study. We also thank Drs. Yasuhiro Saito and Miyuki Shimojo for their assistance in locating FASTQ files required for public sequencing data deposition. The authors used ChatGPT (OpenAI) to assist with language editing and formatting adaptation during manuscript preparation. All AI-assisted text was reviewed and edited by the authors, who take full responsibility for the content of the manuscript. Final English-language editing was performed by American Journal Experts (AJE).

## Author contributions

Conceptualization: J.O., H.K.

Methodology: J.O., H.K., Y.F.

Software: J.O.

Formal analysis: J.O.

Investigation: J.O., M.K., K.F., Y.F.

Data curation: J.O.

Visualization: J.O.

Supervision: H.K.

Funding acquisition: J.O., H.K., A.S.

Project administration: J.O., H.K.

Writing—original draft: J.O.

Writing—review & editing: J.O., H.K., M.K., K.F., Y.F., A.S.

## Data Availability Statement

Processed peptide-count datasets, figure-level input tables, analysis scripts, supplementary tables, and null-model analysis files supporting the findings of this study are available in the GitHub repository at https://github.com/junkookano/constraint_driven_peptide_landscapes and are archived at Zenodo under the version-specific DOI 10.5281/zenodo.20739914.

Raw sequencing reads have been deposited in the DDBJ Sequence Read Archive under BioProject PRJDB42511, with run accessions DRR1062926–DRR1063128 for the *in vivo*-selected datasets and DRR1054851 and DRR1054852 for the same-lot naïve-library datasets. These records are scheduled for release on 2028-06-15 unless released earlier upon publication.

Data supporting the findings of this study are included in the paper and the Supplementary Materials. Reagents used in the exploratory EVGTARY–BPEI polyplex experiment were consumed during the study. The available peptide sequence, reagent information, synthesis details, and experimental conditions are provided in the Methods.

