## Supplementary Figures S1-S22 for "Systemic *in vivo* phage selection reveals compartment-dependent organization of recoverable peptide repertoires"


**This PDF file includes:**

Figs. S1 to S22

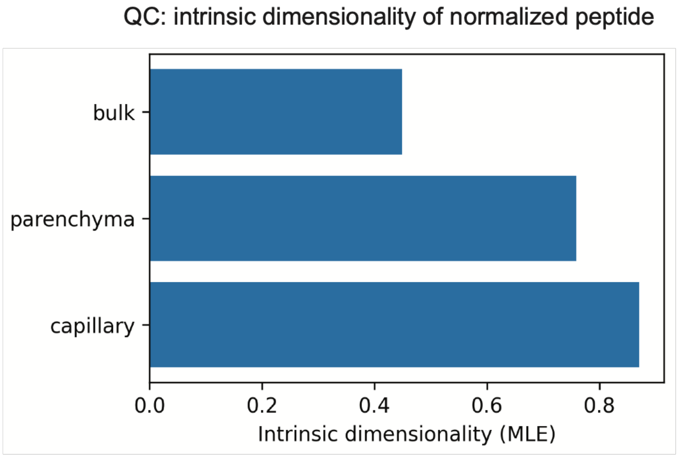


**Fig. S1. Intrinsic dimensionality of the pre-UMAP peptide spaces.**

Intrinsic dimensionality estimates were calculated for the normalized peptide-count spaces before low-dimensional embedding. The estimates provide a QC measure of effective dimensionality across the bulk, capillary, and parenchymal datasets.

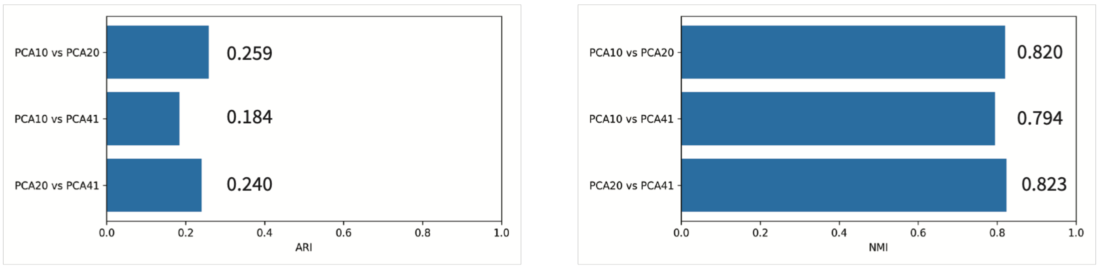


**Fig. S2. Robustness of clustering to PCA dimensionality.**

The clustering concordance was evaluated across PCA dimensionalities (n_pca = 10, 20, and 41; 41 was the maximum feasible value given the 41 organ features). Fine-grained cluster assignments varied modestly, as measured by the adjusted Rand index (ARI ≈ 0.18–0.26), whereas normalized mutual information was high (NMI ≈ 0.79–0.82), supporting robustness of the global clustering structure to PCA dimensionality.

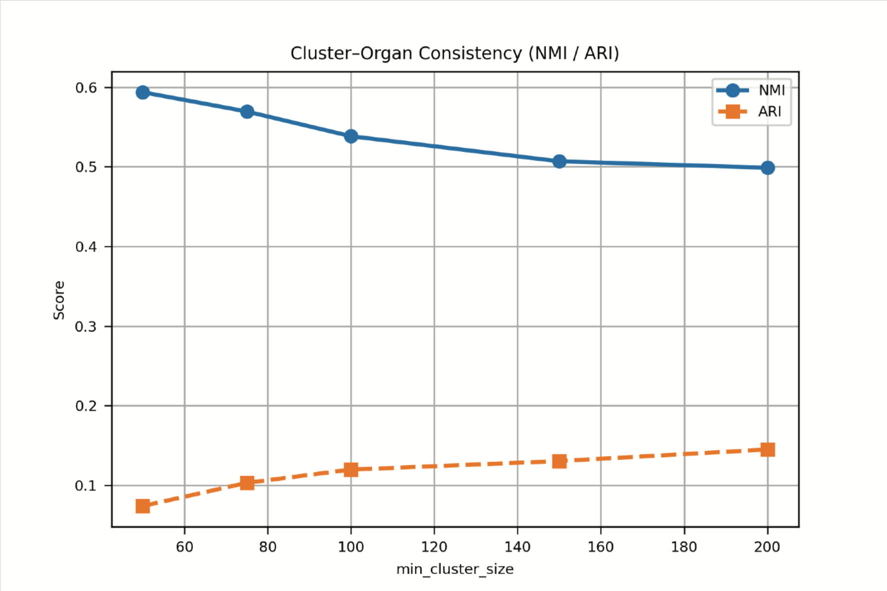


**Fig. S3. Cluster–organ consistency across HDBSCAN parameter settings.**

HDBSCAN cluster assignments generated from the bulk embedding were compared post hoc with organ labels across a range of minimum cluster size (mcs) values. The concordance was quantified using the adjusted Rand index (ARI) and normalized mutual information (NMI). This analysis illustrates the sensitivity of clustering structure to mcs and identifies a parameter regime in which organ-level consistency was maintained without excessive cluster fragmentation. The selected value, mcs = 50, was used for downstream analyses.

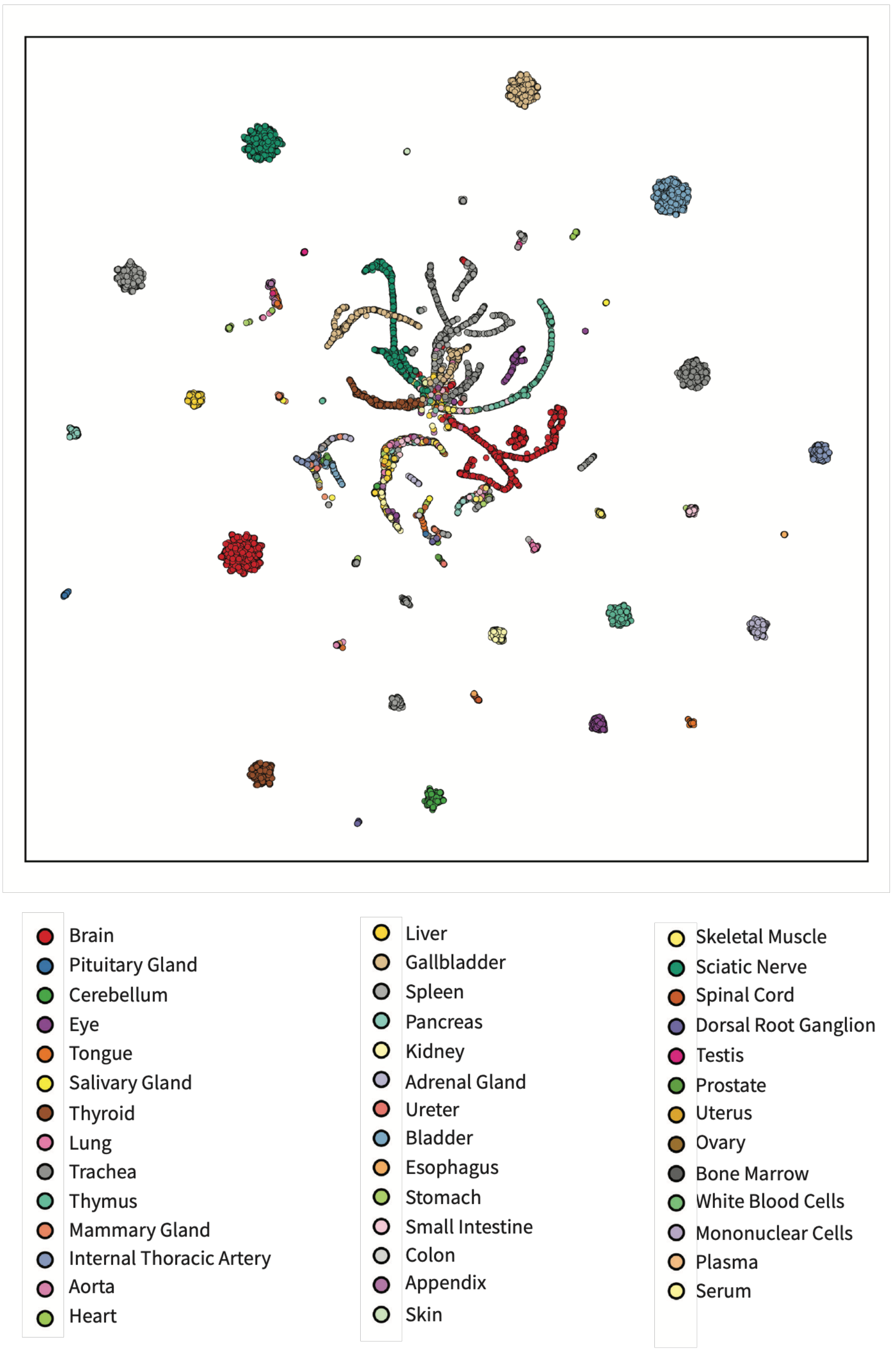


**Fig. S4. Organ-level coloring of the bulk constellation UMAP.**

The UMAP embedding shown in Fig. 1B with physiological-system coloring is shown here with individual organ identities. Organ-level coloring shows finer-grained structure within each physiological system and illustrates the organ-level peptide distributions underlying the system-level constellation pattern.

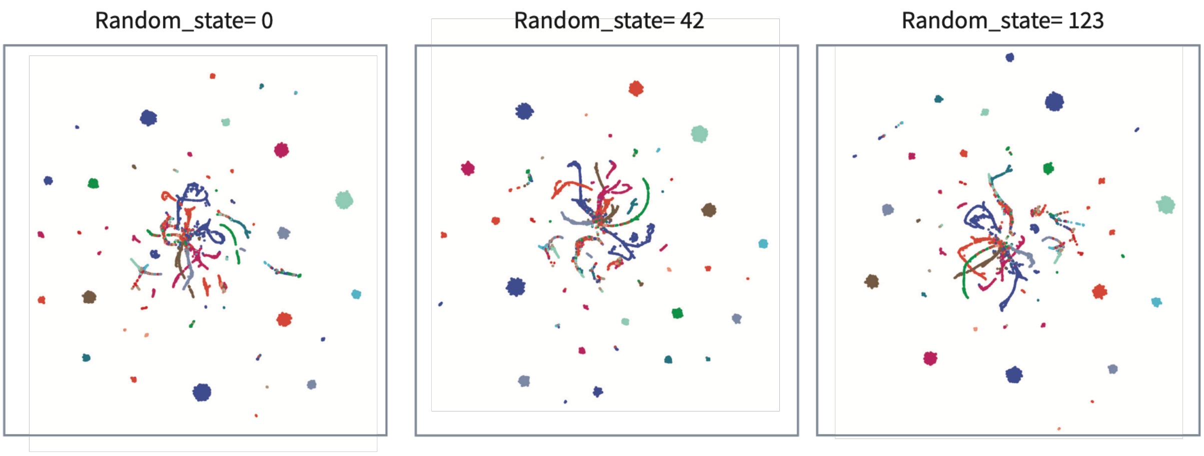


**Fig. S5. Robustness of the bulk system UMAP to random initialization.**

The UMAP embeddings were recomputed by using different random initializations (random_state = 0, 42, 123) with otherwise identical preprocessing procedures and parameters. While the exact layouts differed due to the stochastic nature of the optimization, the global constellation structure, including the shared core and peripheral island organization, was preserved, supporting the robustness of the global embedding structure.

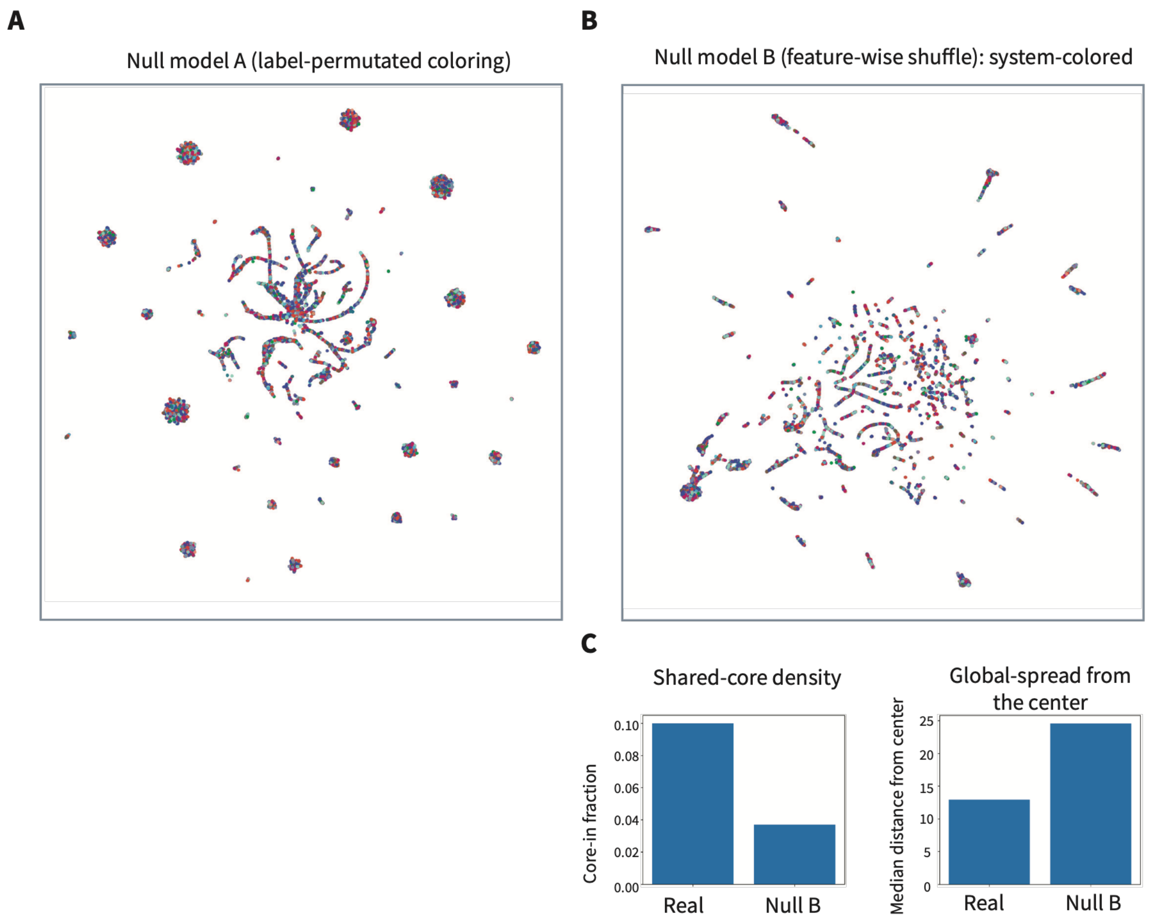


**Fig. S6. Null models for the bulk embedding.**

**A** Label-permutation (color-only) control. The UMAP coordinates were kept fixed while the system labels were randomly permuted. Under permutation, the system colors became spatially intermixed across the embedding, supporting that the coherent system-level structure observed in Fig. 1B was not a consequence of random assignment of system labels to embedding coordinates.

**B** Feature-wise shuffle structure null. Each feature column was independently permuted prior to PCA and UMAP, which disrupted the embedding structure, indicating dependence on the original feature-to-sample correspondences. The system colors are identical to those used in Fig. 1B.

**C** Quantification is shown for the real embedding (Fig. 1B) and the null model B (feature-wise shuffle). The shared-core density was quantified as the fraction of points within a core radius defined by the 10th percentile of distances in the real embedding, and the global spread was quantified as the median distance from the embedding centroid in the real embedding. The global spread was quantified as the median distance from the embedding centroid. Feature-wise shuffle reduced the core-in fraction and increased the median distance, indicating attenuation of the shared-core structure. Similar results were obtained when alternative core percentiles (5–15%) were used.

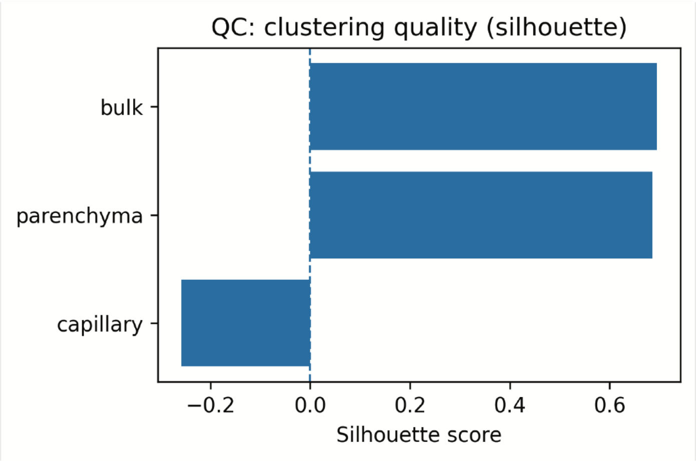


**Fig. S7. QC of the clustering structure (silhouette score).**

Silhouette scores were used as a descriptive QC measure of cluster separability in the UMAP space. Negative values observed in the capillary embedding indicate overlapping neighborhoods and are consistent with a more continuous organization rather than sharply separated clusters.

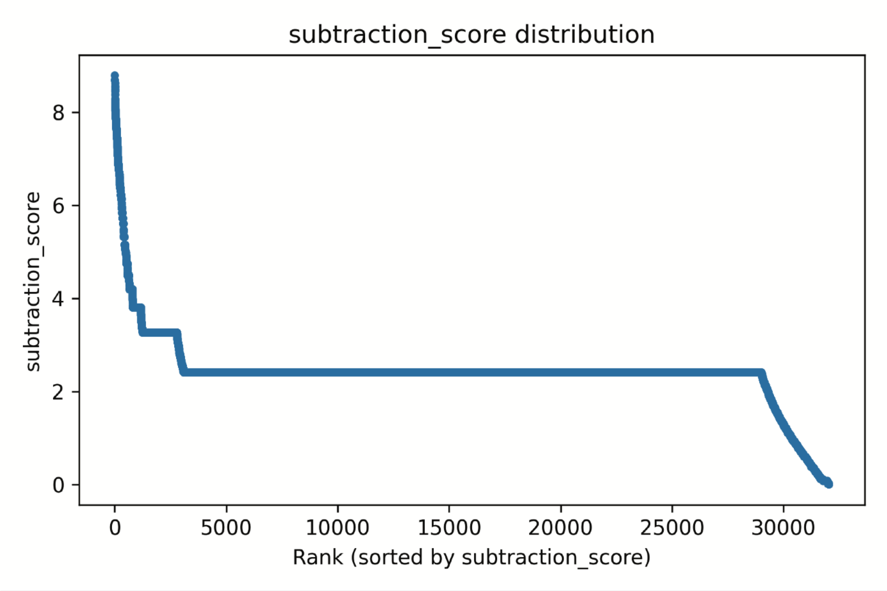


**Fig. S8. Digital subtraction rank curve for the cerebrum-focused bulk analysis.**

Full-rank view of the cerebrum-focused digital subtraction analysis in the bulk dataset. Peptides are ranked by the cerebrum subtraction score, defined as cerebrum enrichment minus the maximum enrichment observed across all other organs. Ranks 1–30,000 are shown to visualize the broader rank distribution beyond the top-ranked region shown in Fig. 2A. The curve shows a stepwise decline with multiple plateau regions.

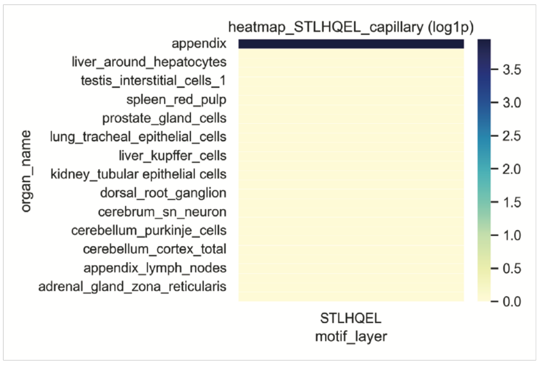


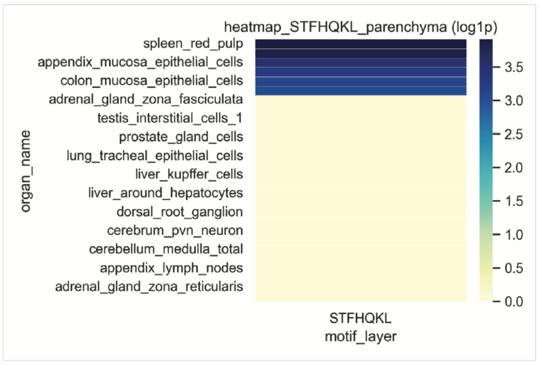


**Fig. S9. Raw read–level enrichment patterns of the medoid sequences identified in Fig. 3A.**

The heatmaps (log1p-transformed raw read counts) show the organ-resolved abundance profiles of the two medoid sequences. STLHQEL, the capillary-anchored medoid, shows highly localized enrichment in the appendix capillary bed (left), whereas STFHQKL, the parenchyma-anchored medoid, shows a broader distribution across multiple organs (right). These results confirm that the medoids correspond to observed enriched peptide sequences with distinct organ distribution profiles.

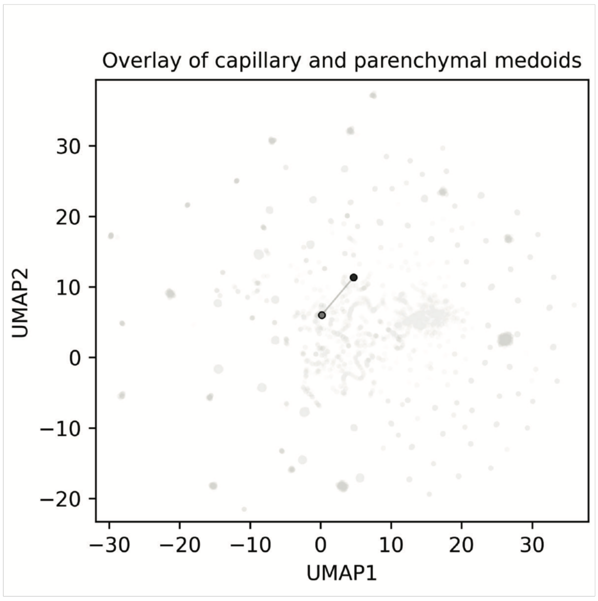


**Fig. S10. Overlay of capillary- and parenchyma-anchored medoids in the shared UMAP embedding.**

The capillary-anchored medoid STLHQEL and parenchyma-anchored medoid STFHQKL are overlaid on the shared UMAP embedding, with STLHQEL shown in light gray and STFHQKL shown in dark gray. This visualization shows their relative positions within the same embedding space and provides a reference for their spatial relationship in the context of the combined capillary–parenchymal peptide distribution.

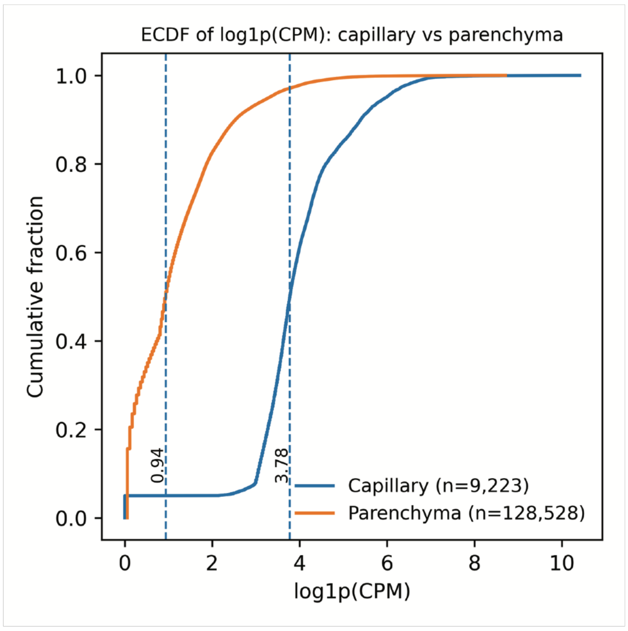


**Fig. S11. Empirical cumulative distribution functions (ECDFs) of the log1p-CPM values for the capillary and parenchyma datasets.**

Empirical cumulative distribution functions (ECDFs) were calculated separately for the capillary (blue) and parenchyma (orange) datasets using log1p-transformed CPM values. The ECDFs show broadly similar distributional shapes, although a global shift between the two datasets is observed. This analysis provides a distributional QC of the normalized values used for visualization and ranking analyses. Dashed vertical lines indicate median log1p-transformed CPM values, 0.94 for capillary and 3.78 for parenchyma. N indicates the number of peptide–sample observations used to construct each ECDF.

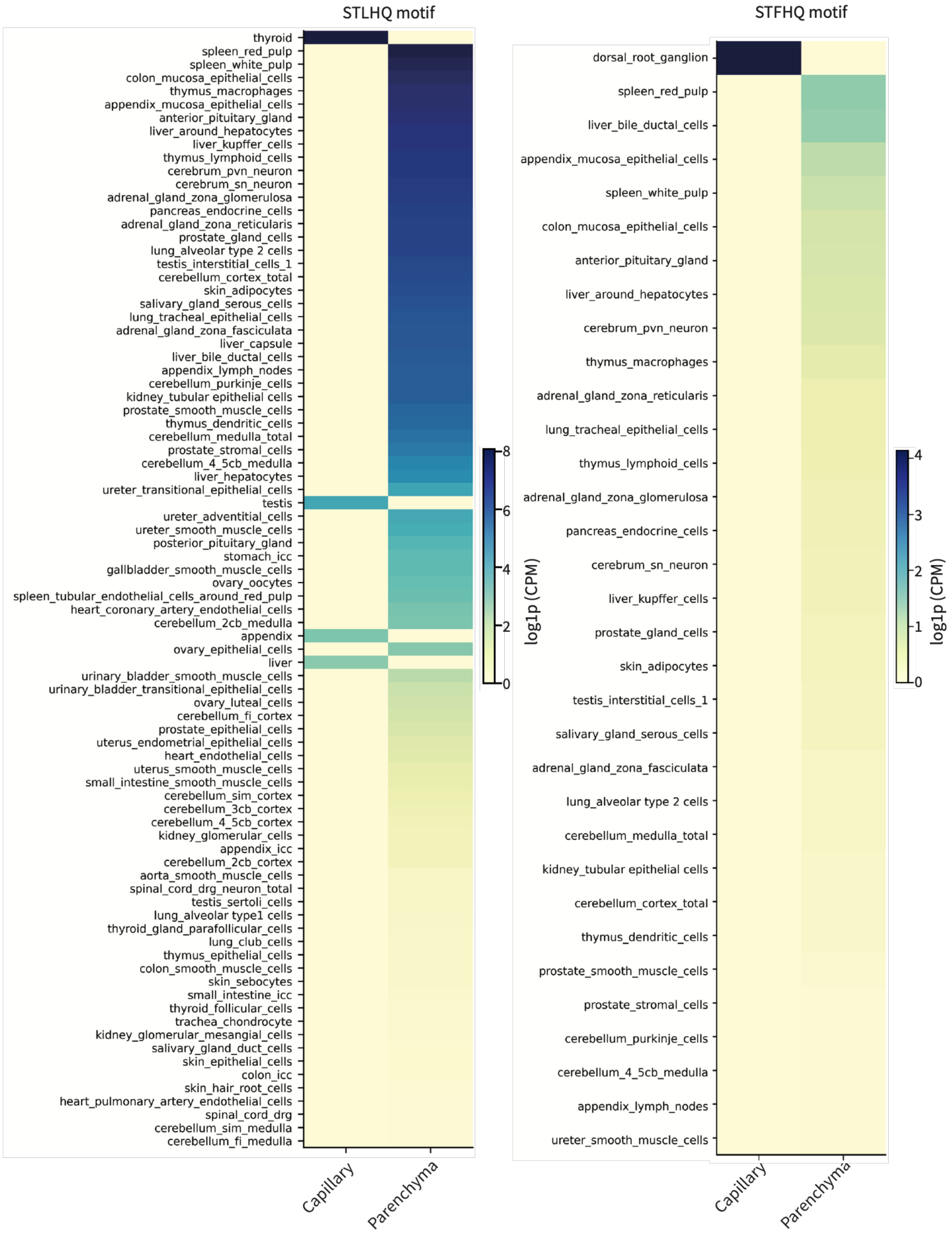


**Fig. S12. Anatomical-site-resolved enrichment heatmaps for the STLHQ- and STFHQ- motifs across the capillary and parenchyma datasets.**

These heatmaps are extended versions of Fig. 3C and show the log1p-CPM–transformed values for each LCM-derived anatomical site, with columns corresponding to the capillary and parenchyma datasets. Anatomical sites are ordered within each motif by their overall ranking based on the log1p-CPM values across both the capillary and parenchyma datasets.
The color scales are independently normalized within each motif family and clipped at the 99th percentile of the nonzero log1p-CPM values.

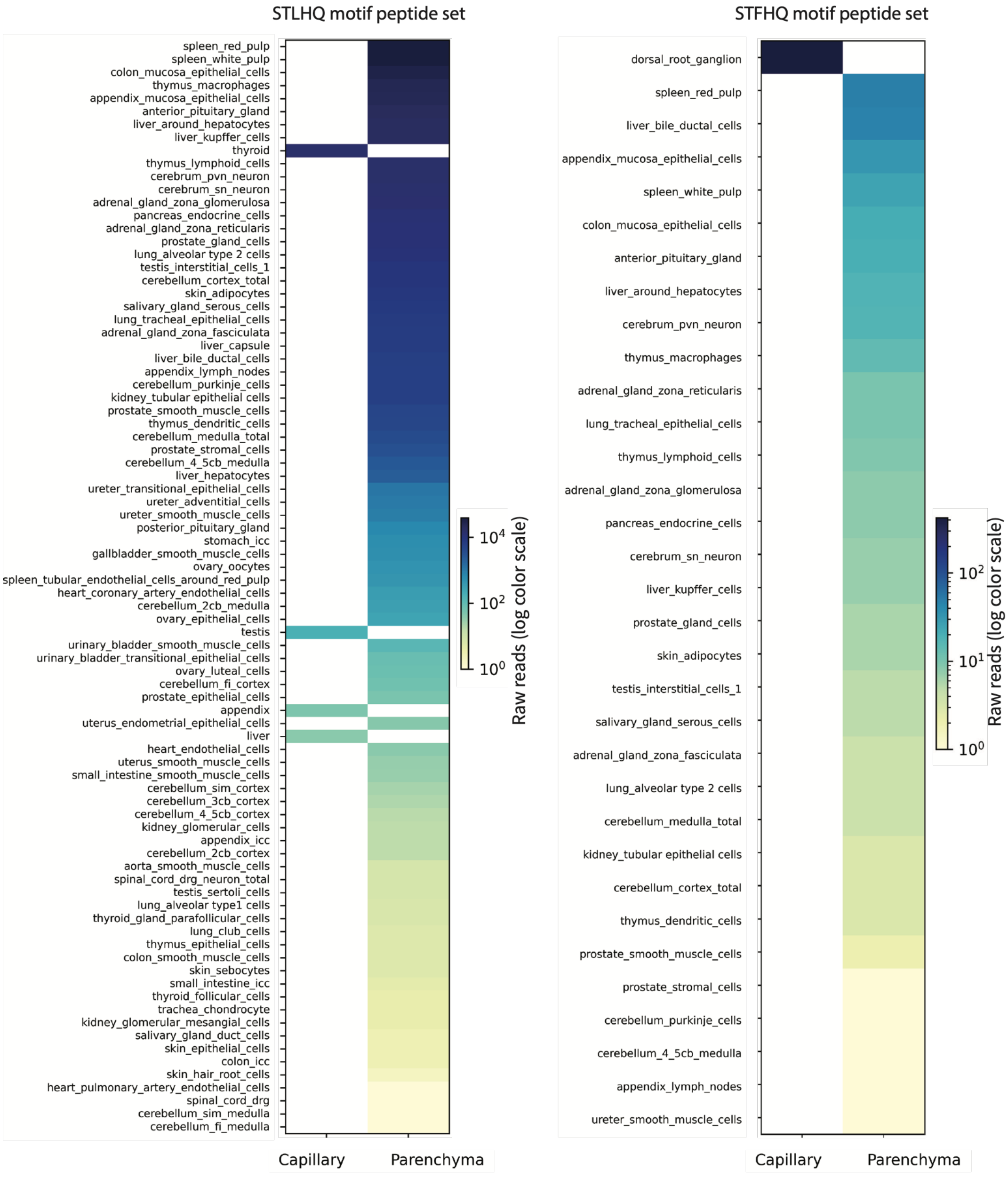


**Fig. S13 | Raw read heatmaps for the STLHQ and STFHQ motif peptide sets across the capillary and parenchyma datasets.**

These heatmaps show raw read counts for each anatomical site, with capillary and parenchyma datasets shown as separate columns and displayed using a logarithmic color scale. The plots provide a raw-read-level comparison with the patterns observed in the log1p-transformed CPM heatmaps. Within each motif, anatomical sites are ordered by the combined raw-read signal across the capillary and parenchyma columns. White cells indicate zero detected reads.

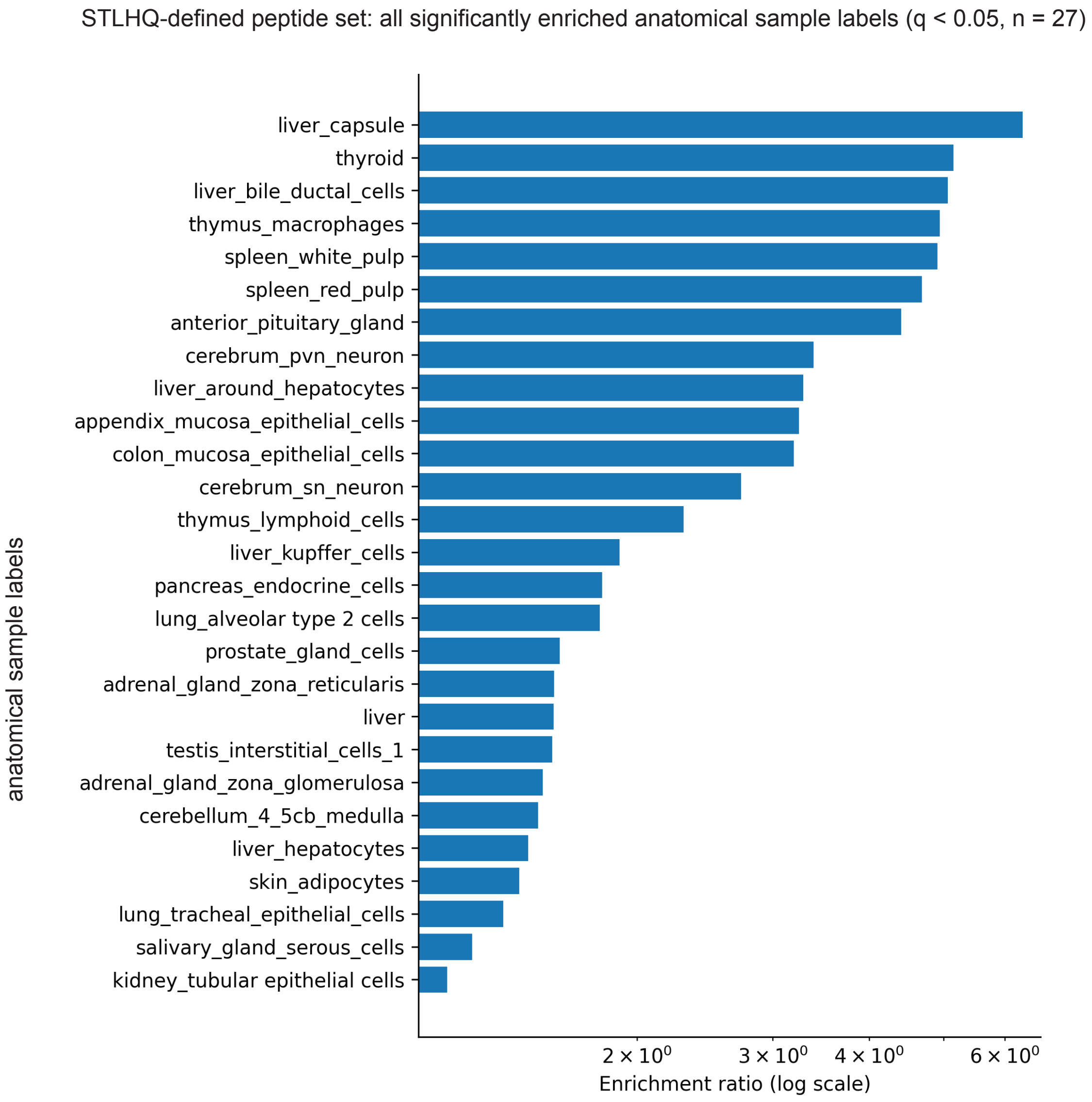


**Fig. S14. Complete enrichment-ratio ranking of anatomical sample labels for the STLHQ-defined peptide set.**

This figure extends Fig. 3E by showing the complete set of anatomical sample labels that met the significance threshold for enrichment of the STLHQ-defined peptide set ($q < 0.05$, $n = 27$). Bars represent anatomical-sites-wise enrichment ratios calculated for each anatomical site, and the x-axis is shown on a logarithmic scale.

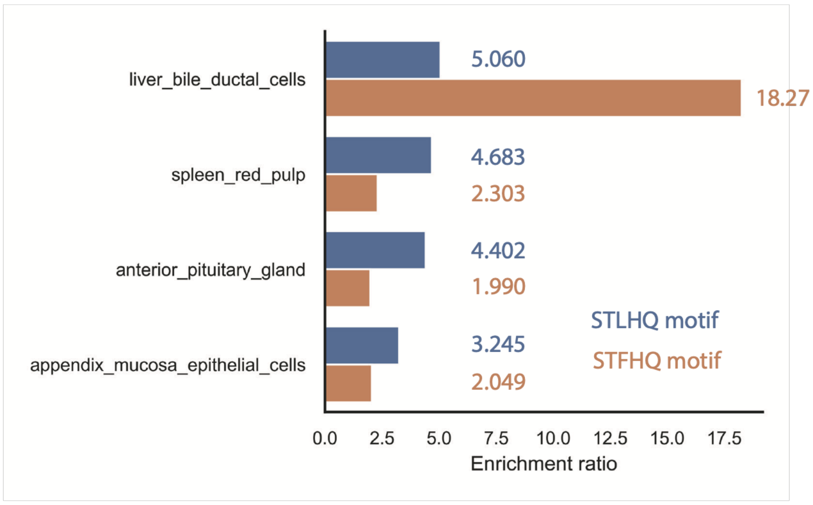


**Fig. S15. Individual enrichment ratios underlying the motif convergence score for anatomical sample labels with concordant enrichment.**

The bar plots show the individual enrichment ratios of the STLHQ-defined peptide set (blue) and STFHQ-defined peptide set (orange) for each anatomical sample label shown in Fig. 3E, right. These values were used to calculate the motif convergence score. Numerical values next to each bar indicate the corresponding enrichment ratios.

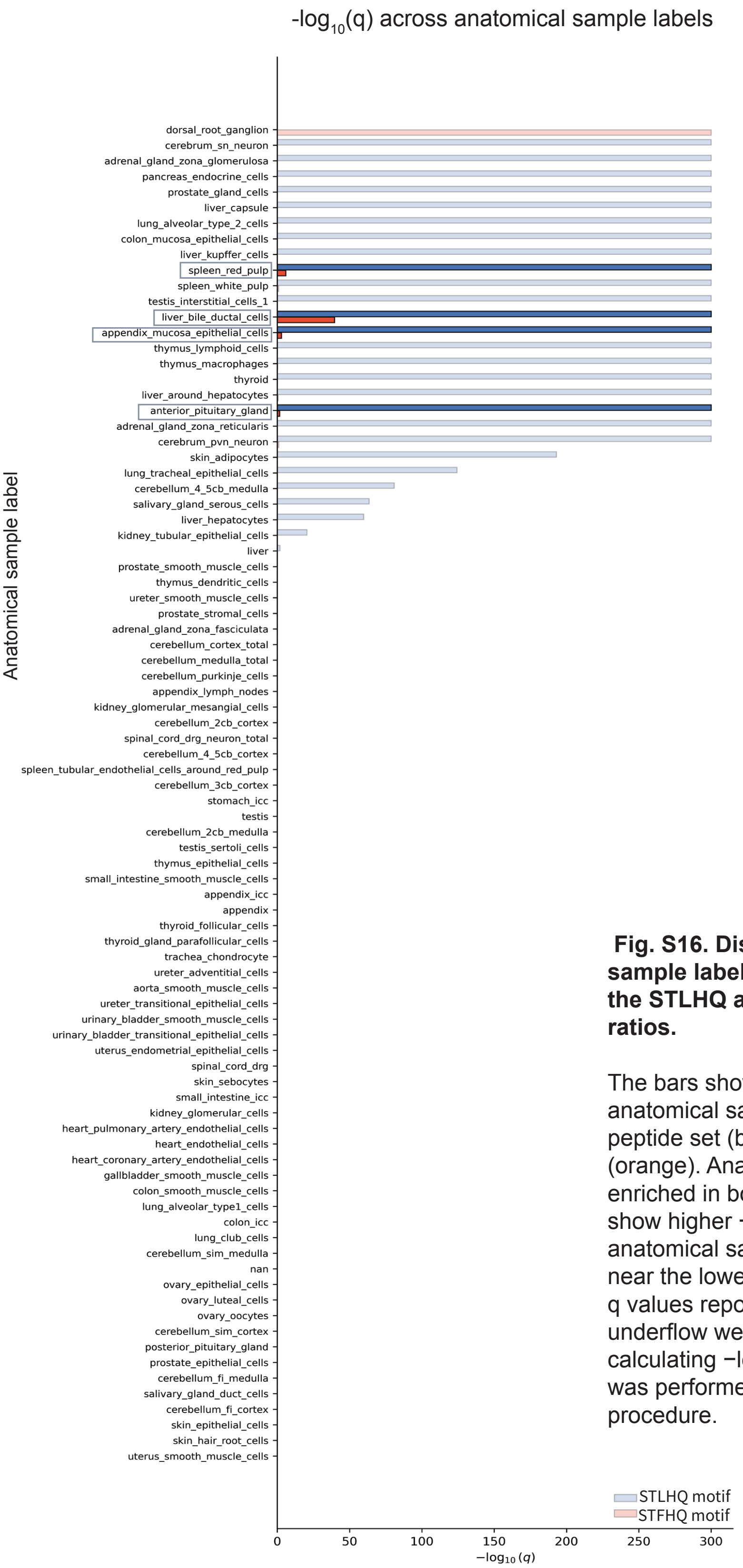


**Fig. S16. Distribution across anatomical sample labels of the statistical significance for the STLHQ and STFHQ motif enrichment ratios.**

The bars show −log10(q) values for each anatomical sample label for the STLHQ-defined peptide set (blue) and STFHQ-defined peptide set (orange). Anatomical sample labels significantly enriched in both motif analyses, outlined in gray, show higher −log10(q) values than most other anatomical sample labels, which are distributed near the lower end of the scale. For visualization, q values reported as 0 because of numerical underflow were clipped to $1 \times 10^{-300}$ before calculating −log10(q). Multiple-testing correction was performed using the Benjamini–Hochberg procedure.

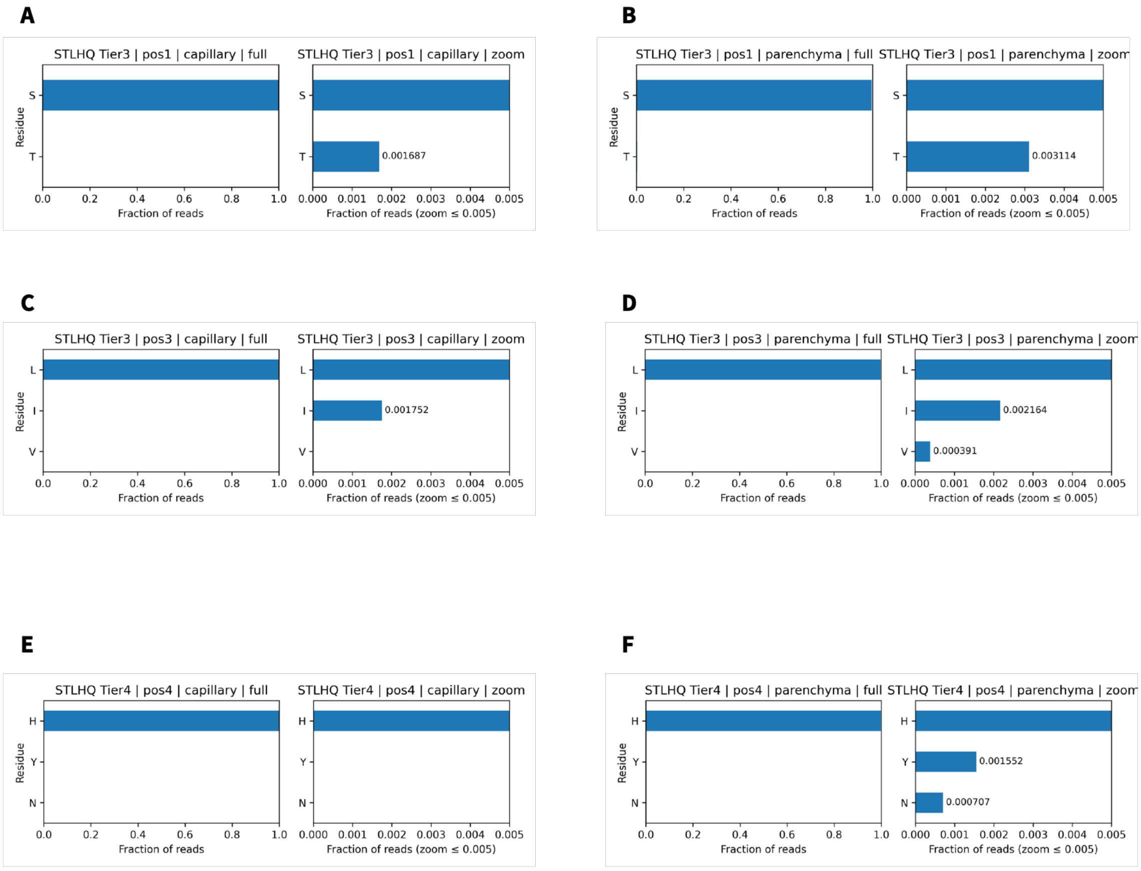


**Fig. S17. Read-weighted residue fractions under tiered perturbation of the STLHQ motif core.**

Read-weighted horizontal bar plots show residue frequencies at selected positions in the STLHQ-defined peptide sets under predefined perturbation tiers, shown separately for the capillary and parenchyma compartments. For each condition, the left panel shows the full-scale fraction of reads, 0–1, and the right panel shows a zoomed-in view, 0–0.005, to visualize low-frequency deviations that are not readily apparent in sequence-logo representations.

Tier 3 core relaxation, ^[ST][TL][IVL]HQ..$: residue frequencies at position 1 are shown for the capillary (A) and parenchyma (B) compartments, and residue frequencies at position 3 are shown for the capillary (C) and parenchyma (D) compartments. Across compartments, the dominant residues remained highly enriched, with substitutions constrained to sub-percent levels.

Tier 4 H-site test, ^STL[HNY]Q..$: residue frequencies at position 4 are shown for the capillary (E) and parenchyma (F) compartments. Capillary sequences remained fixed at H, whereas parenchyma sequences showed rare H→Y/N substitutions at low frequency.

Sequence-logo representations for Tier 3 and Tier 4 did not show readily discernible differences from the corresponding lower-tier logos; therefore, read-weighted bar plots are shown to resolve low-frequency deviations at the selected positions.

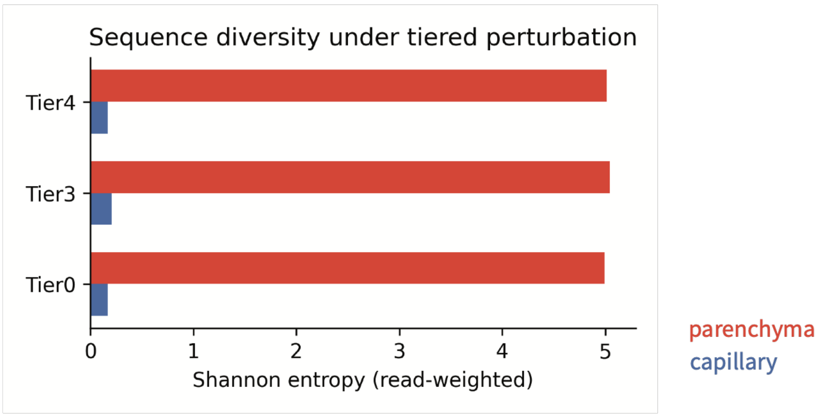


**Fig. S18 | Shannon entropy in the capillary and parenchyma datasets across tiers.**

Shannon entropy was calculated as a read-weighted sequence-level measure of diversity for the STLHQ-defined peptide sets in the capillary and parenchyma datasets. Across Tier 0, Tier 3, and Tier 4, entropy values remained approximately 5 in the parenchyma dataset and 0.17 in the capillary dataset, despite the expanded perturbation definitions. The stable separation is consistent with greater read-weighted sequence diversity in the parenchyma-associated STLHQ-defined peptide set than in the capillary-associated set.

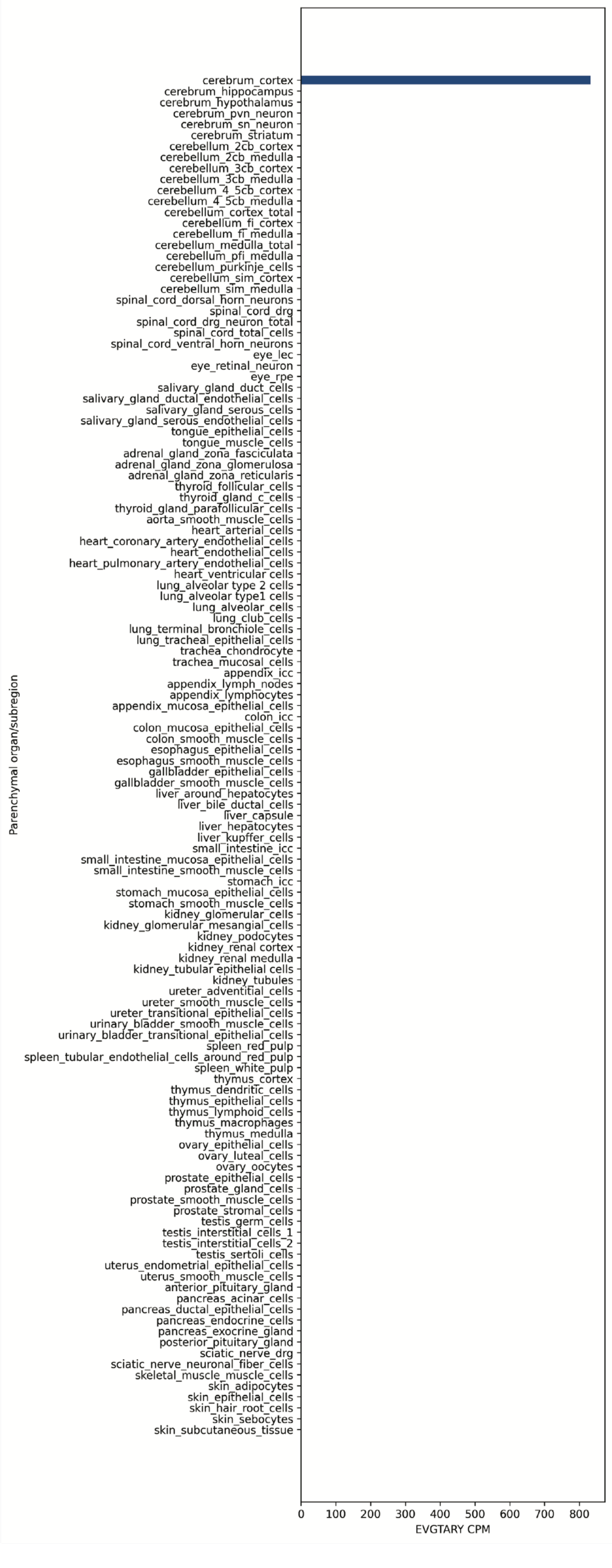


**Fig. S19. Anatomical distribution of EVGTARY in the parenchyma-associated dataset (CPM-normalized).**

EVGTARY was detected in cerebral cortex-derived parenchymal samples but was not detected in other parenchymal samples within the screening dataset, supporting its prioritization as a brain-associated candidate for exploratory proof-of-concept testing.

**A** **B**

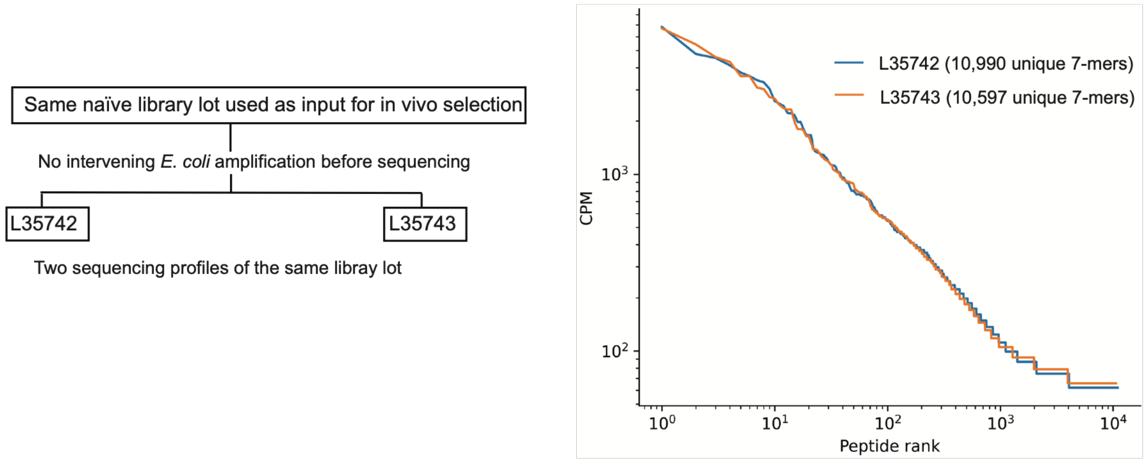


**C**

| Query type | Feature | L35742 matched reads | L35743 matched reads |
|---|---|---|---|
| Exact peptide | STLHQEL | 0 | 0 |
| Exact peptide | STFHQKL | 0 | 0 |
| Exact peptide | EVGTARY | 0 | 0 |
| Motif | STLHQxx | 0 | 0 |
| Motif | STFHQxx | 0 | 0 |

**Fig. S20. Same-lot naïve phage library background profiling.**

**A)** Provenance of the naïve-library datasets. L35742 and L35743 were generated as two sequencing profiles of the same naïve phage library lot used as the input for the mouse in vivo selection experiments. No intervening E. coli amplification of the phage library was performed before sequencing. The two datasets therefore represent profiles of the same input library lot rather than independent library lots or biological replicates.

**B)** Rank–abundance distributions of valid 7-mer peptide sequences detected in L35742 and L35743. CPM values and peptide ranks were calculated separately for each profile without pooling. The profiles contained 10,990 and 10,597 detected unique 7-mer sequences, respectively. Both axes are shown on logarithmic scales.

**C)** Targeted queries for the exact peptide sequences STLHQEL, STFHQKL, and EVGTARY and for the STLHQxx and STFHQxx motifs in the two naïve-library profiles. The motif queries identified valid 7-mer sequences matching the anchored patterns ^STLHQ..$ and ^STFHQ..$, respectively, where each x represents any standard amino acid. None of the queried exact peptides or motif-matching sequences was detected in either L35742 or L35743.

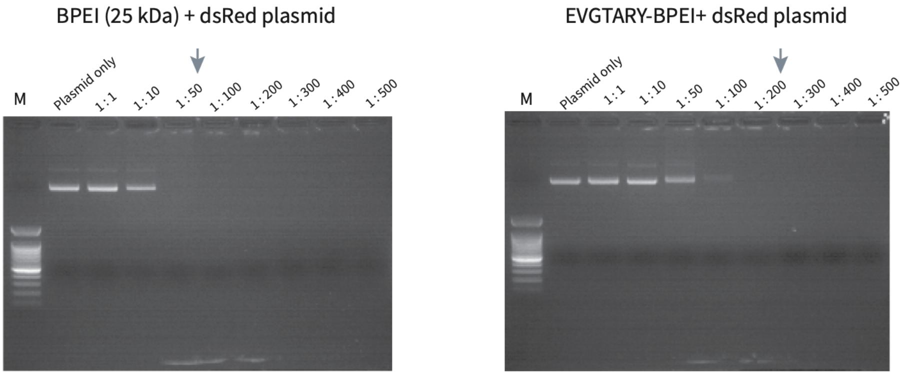


**Fig. S21. Gel retardation assay showing the charge-mediated association of dsRed plasmid with BPEI-based polyplexes.**

Agarose gel electrophoresis of dsRed plasmid mixed with BPEI (left) or EVGTARY-modified BPEI (right) at increasing charge-neutralization-based polymer-to-DNA equivalent ratios. Reduced plasmid mobility at higher ratios is consistent with charge-mediated plasmid–polymer complexation and polyplex formation. Arrows indicate the formulation conditions selected for the in vivo experiments: BPEI, 1:50; EVGTARY–BPEI, 1:200. M, DNA ladder.

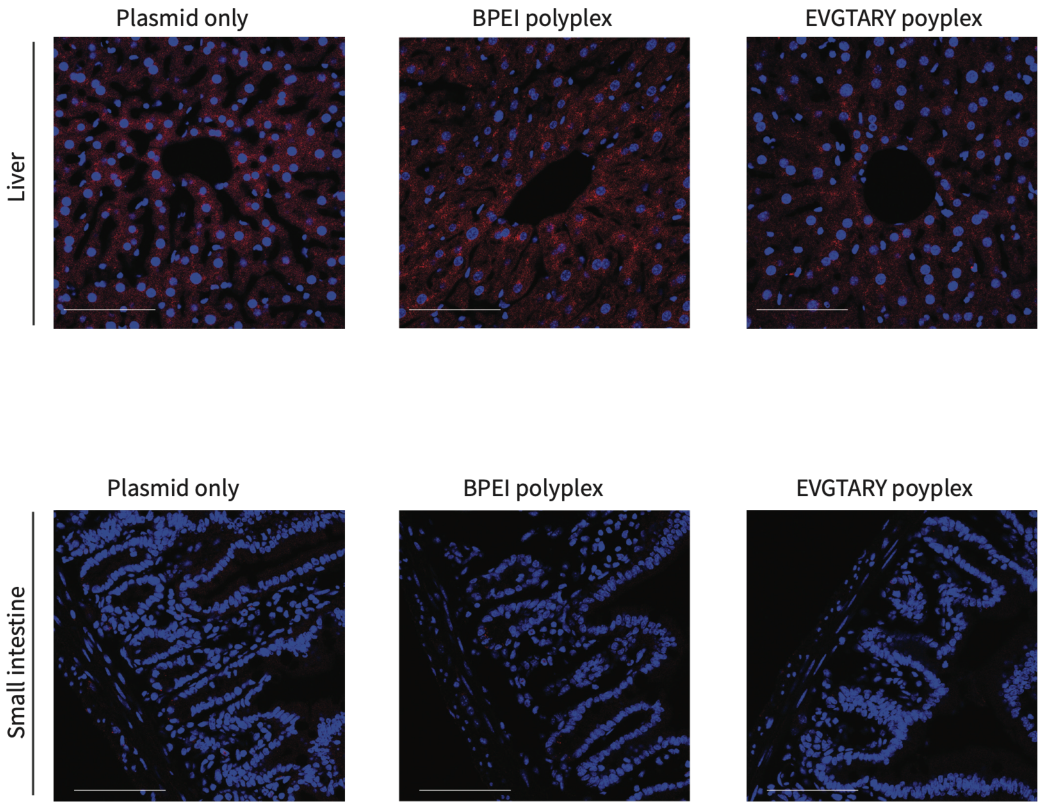


**Fig. S22. Liver and intestinal sections following systemic administration.**

Upper panels, representative liver sections from mice injected with plasmid only, BPEI polyplex, or EVGTARY polyplex show comparably low-level dsRed signals across conditions.

Lower panels, representative intestinal sections from mice injected with plasmid only, BPEI polyplex, or EVGTARY polyplex show no apparent dsRed signal. Scale bars, 100 µm.

Images were acquired and displayed using identical microscope settings and contrast/lookup table (LUT) scaling as described for Fig. 4C.
