## Supplementary Tables S1-S5 for "Systemic *in vivo* phage selection reveals compartment-dependent organization of recoverable peptide repertoires"

#### Table S1. Organ classification used for UMAP visualization in Figure 1B

| Organ | System |
|---|---|
| Cerebrum | Nervous |
| Cerebellum | Nervous |
| Spinal cord | Nervous |
| Dorsal root ganglion | Nervous |
| Sciatic nerve | Nervous |
| Pituitary gland | Nervous |
| Heart | Vascular |
| Internal thoracic artery | Vascular |
| Aorta | Vascular |
| Spleen | Immune |
| Thymus | Immune |
| Appendix | Immune |
| Liver | Digestive |
| Gallbladder | Digestive |
| Pancreas | Digestive |
| Esophagus | Digestive |
| Stomach | Digestive |
| Small intestine | Digestive |
| Colon | Digestive |
| Lung | Respiratory |
| Trachea | Respiratory |
| Skin | Skin/Musculo |
| Skeletal muscle | Skin/Musculo |
| Kidney | Urinary |
| Ureter | Urinary |
| Bladder | Urinary |
| Thyroid | Endocrine |
| Adrenal gland | Endocrine |
| Salivary gland | Exocrine |
| Mammary gland | Exocrine |
| Testis | Genital |
| Prostate | Genital |
| Uterus | Genital |
| Ovary | Genital |
| Eye | Sensory |

| Organ | System |
| --- | --- |
| Tongue | Sensory |
| Bone marrow | Blood/Hematopoietic |
| White blood cells | Blood/Hematopoietic |
| Mononuclear cells | Blood/Hematopoietic |
| Plasma | Blood/Hematopoietic |
| Serum | Blood/Hematopoietic |

#### Table S2. Annotation of organ and anatomical subregions corresponding to the plotted symbols used in Fig. 1D

| Label | Plotting symbol | Organ | Subregion |
|---|---|---|---|
| a | circle | cerebrum | cerebrum_cortex |
| b | square | cerebrum | cerebrum_hippocampus |
| c | triangle (up) | cerebrum | cerebrum_hypothalamus |
| d | triangle (down) | cerebrum | cerebrum_sn_neuron |
| e | diamond | cerebrum | cerebrum_pvn_neuron |
| f | plus | cerebrum | cerebrum_striatum |
| a | circle | pituitary gland | anterior_pituitary_gland |
| b | square | pituitary gland | posterior_pituitary_gland |
| a | circle | cerebellum | cerebellum_2cb_cortex |
| b | square | cerebellum | cerebellum_3cb_cortex |
| c | triangle (up) | cerebellum | cerebellum_4_5cb_cortex |
| d | triangle (down) | cerebellum | cerebellum_fi_cortex |
| e | diamond | cerebellum | cerebellum_pfi_medulla |
| f | plus | cerebellum | cerebellum_sim_cortex |
| g | cross | cerebellum | cerebellum_purkinje_cells |
| h | star | cerebellum | cerebellum_2cb_medulla |
| i | triangle (left) | cerebellum | cerebellum_3cb_medulla |
| j | triangle (right) | cerebellum | cerebellum_4_5cb_medulla |
| k | pentagon | cerebellum | cerebellum_fi_medulla |
| l | hexagon | cerebellum | cerebellum_cortex_total |
| m | octagon | cerebellum | cerebellum_sim_medulla |
| n | thin diamond | cerebellum | cerebellum_medulla_total |
| a | circle | eye | eye_retinal_neuron |
| b | square | eye | eye_rpe |
| c | triangle (up) | eye | eye_lec |
| a | circle | tongue | tongue_muscle_cells |
| b | square | tongue | tongue_epithelial_cells |
| a | circle | salivary gland | salivary_gland_duct_cells |
| b | square | salivary gland | salivary_gland_ductal_endothelial_cells |
| c | triangle (up) | salivary gland | salivary_gland_serous_cells |
| d | triangle (down) | salivary gland | salivary_gland_serous_endothelial_cells |
| a | circle | thyroid gland | thyroid_gland_c_cells |
| b | square | thyroid gland | thyroid_follicular_cells |
| c | triangle (up) | thyroid gland | thyroid_gland_parafollicular_cells |
| a | circle | lung | lung_club_cells |

| Label | Plotting symbol | Organ | Subregion |
|---|---|---|---|
| b | square | lung | lung_alveolar_cells |
| c | triangle (up) | lung | lung_tracheal_epithelial_cells |
| d | triangle (down) | lung | lung_terminal_bronchiole_cells |
| e | diamond | lung | lung_alveolar type1 cells |
| f | plus | lung | lung_alveolar type 2 cells |
| a | circle | trachea | trachea_chondrocyte |
| b | square | trachea | trachea_mucosal_cells |
| a | circle | thymus | thymus_cortex |
| b | square | thymus | thymus_dendritic_cells |
| c | triangle (up) | thymus | thymus_epithelial_cells |
| d | triangle (down) | thymus | thymus_lymphoid_cells |
| e | diamond | thymus | thymus_macrophages |
| f | plus | thymus | thymus_medulla |
| a | circle | aorta | aorta_smooth_muscle_cells |
| a | circle | heart | heart_coronary_artery_endothelial_cells |
| b | square | heart | heart_pulmonary_artery_endothelial_cells |
| c | triangle (up) | heart | heart_endothelial_cells |
| d | triangle (down) | heart | heart_atrial_cells |
| e | diamond | heart | heart_ventricular cells |
| a | circle | liver | liver_around_hepatocytes |
| b | square | liver | liver_bile_ductal_cells |
| c | triangle (up) | liver | liver_hepatocytes |
| d | triangle (down) | liver | liver_capsule |
| e | diamond | liver | liver_kupffer_cells |
| a | circle | gallbladder | gallbladder_epithelial_cells |
| b | square | gallbladder | gallbladder_smooth_muscle_cells |
| a | circle | spleen | spleen_white_pulp |
| b | square | spleen | spleen_tubular_endothelial_cells_around_red_pulp |
| c | triangle (up) | spleen | spleen_red_pulp |
| a | circle | pancreas | pancreas_ductal_epithelial_cells |
| b | square | pancreas | pancreas_endocrine_cells |
| c | triangle (up) | pancreas | pancreas_exocrine_gland |
| d | triangle (down) | pancreas | pancreas_acinar_cells |
| a | circle | kidney | kidney_glomerular_cells |
| b | square | kidney | kidney_glomerular_mesangial_cells |
| c | triangle (up) | kidney | kidney_tubular epithelial_cells |
| d | triangle (down) | kidney | kidney_tubules |

| Label | Plotting symbol | Organ | Subregion |
|---|---|---|---|
| e | diamond | kidney | kidney_podocytes |
| f | plus | kidney | kidney_renal cortex |
| g | cross | kidney | kidney_renal medulla |
| a | circle | adrenal gland | adrenal_gland_zona_glomerulosa |
| b | square | adrenal gland | adrenal_gland_zona_reticularis |
| c | triangle (up) | adrenal gland | adrenal_gland_zona_fasciculata |
| a | circle | ureter | ureter_adventitial_cells |
| b | square | ureter | ureter_transitional_epithelial_cells |
| c | triangle (up) | ureter | ureter_smooth_muscle_cells |
| a | circle | urinary bladder | urinary_bladder_transitional_epithelial_cells |
| b | square | urinary bladder | urinary_bladder_smooth_muscle_cells |
| a | circle | esophagus | esophagus_epithelial_cells |
| b | square | esophagus | esophagus_smooth_muscle_cells |
| a | circle | stomach | stomach_icc |
| b | square | stomach | stomach_mucosa_epithelial_cells |
| c | triangle (up) | stomach | stomach_smooth_muscle_cells |
| a | circle | small intestine | small_intestine_icc |
| b | square | small intestine | small_intestine_mucosa_epithelial_cells |
| c | triangle (up) | small intestine | small_intestine_smooth_muscle_cells |
| a | circle | colon | colon_icc |
| b | square | colon | colon_mucosa_epithelial_cells |
| c | triangle (up) | colon | colon_smooth_muscle_cells |
| a | circle | appendix | appendix_icc |
| b | square | appendix | appendix_mucosa_epithelial_cells |
| c | triangle (up) | appendix | appendix_lymphocytes |
| d | triangle (down) | appendix | appendix_lymph_nodes |
| e | diamond | appendix | appendix_smooth_muscle_cells |
| a | circle | skeletal muscle | skeletal_muscle_muscle_cells |
| a | circle | skin | skin_adipocytes |
| b | square | skin | skin_subcutaneous_tissue |
| c | triangle (up) | skin | skin_epithelial_cells |
| d | triangle (down) | skin | skin_hair_root_cells |
| e | diamond | skin | skin_sebocytes |
| a | circle | sciatic nerve | sciatic_nerve_drg |
| b | square | sciatic nerve | sciatic_nerve_neuronal_fiber_cells |
| a | circle | spinal cord | spinal_cord_total_cells |
| b | square | spinal cord | spinal_cord_dorsal_horn_neurons |

| Label | Plotting symbol | Organ | Subregion |
|---|---|---|---|
| c | triangle (up) | spinal cord | spinal_cord_ventral_horn_neurons |
| a | circle | Dorsal root ganglion | spinal_cord_drg_neuron_total |
| b | square | Dorsal root ganglion | spinal_cord_drg |
| a | circle | testis | testis_interstitial_cells_1 |
| b | square | testis | testis_interstitial_cells_2 |
| c | triangle (up) | testis | testis_germ_cells |
| d | triangle (down) | testis | testis_sertoli_cells |
| a | circle | prostate | prostate_epithelial_cells |
| b | square | prostate | prostate_stromal_cells |
| c | triangle (up) | prostate | prostate_gland_cells |
| d | triangle (down) | prostate | prostate_smooth_muscle_cells |
| a | circle | uterus | uterus_endometrial_epithelial_cells |
| b | square | uterus | uterus_smooth_muscle_cells |
| a | circle | ovary | ovary_oocytes |
| b | square | ovary | ovary_luteal_cells |
| c | triangle (up) | ovary | ovary_epithelial_cells |

Letters (a–n) correspond to the labels indicated in the UMAP visualization.

#### Table S3. Top 100 peptides ranked by the cerebrum subtraction score

| Rank | Peptide sequence | subtraction_score | E_cerebrum | E_liver | E_skeletal_muscle |
|---|---|---|---|---|---|
| 1 | DFGKAHS | 8.788403738 | 8.788403738 | 0 | 0 |
| 2 | GTSNTHG | 8.701144813 | 8.701144813 | 0 | 0 |
| 3 | NALPSKT | 8.686074407 | 8.686074407 | 0 | 0 |
| 4 | LTIGRNN | 8.624167397 | 8.624167397 | 0 | 0 |
| 5 | NQSPYSF | 8.559484348 | 8.559484348 | 0 | 0 |
| 6 | TGRTNAN | 8.542850071 | 8.542850071 | 0 | 0 |
| 7 | RGSTAFH | 8.491764559 | 8.491764559 | 0 | 0 |
| 8 | DKHYVNQ | 8.474326006 | 8.474326006 | 0 | 0 |
| 9 | DRNVLPT | 8.456674084 | 8.456674084 | 0 | 0 |
| 10 | VSKSSTS | 8.383823925 | 8.383823925 | 0 | 0 |
| 11 | LPNKHGS | 8.287262292 | 8.287262292 | 0 | 0 |
| 12 | SNNNEAA | 8.267149451 | 8.267149451 | 0 | 0 |
| 13 | SEKPNSN | 8.267149451 | 8.267149451 | 0 | 0 |
| 14 | HHDHPQL | 8.226062519 | 8.226062519 | 0 | 0 |
| 15 | DKSASSS | 8.205071757 | 8.205071757 | 0 | 0 |
| 16 | ESQLKKS | 8.183771071 | 8.183771071 | 0 | 0 |
| 17 | TSPKMLH | 8.162151173 | 8.162151173 | 0 | 0 |
| 18 | SRNMATM | 8.140202348 | 8.140202348 | 0 | 0 |
| 19 | SKGSALQ | 8.117914433 | 8.117914433 | 0 | 0 |
| 20 | PSTHTHA | 8.117914433 | 8.117914433 | 0 | 0 |
| 21 | SHRNTLV | 8.095276787 | 8.095276787 | 0 | 0 |
| 22 | DVNTTKH | 8.072278258 | 8.072278258 | 0 | 0 |
| 23 | KNALKPQ | 8.072278258 | 8.072278258 | 0 | 0 |
| 24 | MSSNPTR | 8.048907153 | 8.048907153 | 0 | 0 |
| 25 | TTARYPL | 8.048907153 | 8.048907153 | 0 | 0 |
| 26 | WNGVINT | 8.048907153 | 8.048907153 | 0 | 0 |
| 27 | YWKNSMA | 8.025151203 | 8.025151203 | 0 | 0 |
| 28 | SNDVLAW | 8.025151203 | 8.025151203 | 0 | 0 |
| 29 | VNDSAKT | 8.025151203 | 8.025151203 | 0 | 0 |
| 30 | HNSRYQE | 8.000997521 | 8.000997521 | 0 | 0 |
| 31 | SNKINKG | 8.000997521 | 8.000997521 | 0 | 0 |
| 32 | HLHQKYL | 7.97643256 | 7.97643256 | 0 | 0 |
| 33 | NSMSNRV | 7.97643256 | 7.97643256 | 0 | 0 |
| 34 | TLKSGQR | 7.97643256 | 7.97643256 | 0 | 0 |
| 35 | KSIMNPH | 7.97643256 | 7.97643256 | 0 | 0 |

| Rank | Peptide sequence | subtraction_score | E_cerebrum | E_liver | E_skeletal_muscle |
|---|---|---|---|---|---|
| 36 | THGRSSL | 7.951442073 | 7.951442073 | 0 | 0 |
| 37 | VQELMLA | 7.951442073 | 7.951442073 | 0 | 0 |
| 38 | RGGPHQS | 7.951442073 | 7.951442073 | 0 | 0 |
| 39 | DRHYAPF | 7.951442073 | 7.951442073 | 0 | 0 |
| 40 | KTTKASL | 7.926011056 | 7.926011056 | 0 | 0 |
| 41 | DTSVDRK | 7.926011056 | 7.926011056 | 0 | 0 |
| 42 | PIHGRSS | 7.926011056 | 7.926011056 | 0 | 0 |
| 43 | LPLHHSR | 7.926011056 | 7.926011056 | 0 | 0 |
| 44 | MSSSNAF | 7.9001237 | 7.9001237 | 0 | 0 |
| 45 | HVRFPNQ | 7.9001237 | 7.9001237 | 0 | 0 |
| 46 | QPRGQNS | 7.873763327 | 7.873763327 | 0 | 0 |
| 47 | DTRTAAQ | 7.873763327 | 7.873763327 | 0 | 0 |
| 48 | NYSQHYN | 7.846912328 | 7.846912328 | 0 | 0 |
| 49 | TVNNDNS | 7.846912328 | 7.846912328 | 0 | 0 |
| 50 | KNTTEQL | 7.819552095 | 7.819552095 | 0 | 0 |
| 51 | YQRSNNV | 7.819552095 | 7.819552095 | 0 | 0 |
| 52 | TWYSTLA | 7.819552095 | 7.819552095 | 0 | 0 |
| 53 | NRQDLMY | 7.791662936 | 7.791662936 | 0 | 0 |
| 54 | TTHPKDQ | 7.791662936 | 7.791662936 | 0 | 0 |
| 55 | VELGLRS | 7.791662936 | 7.791662936 | 0 | 0 |
| 56 | HWSTSLS | 7.791662936 | 7.791662936 | 0 | 0 |
| 57 | DRTDTEG | 7.763223999 | 7.763223999 | 0 | 0 |
| 58 | QPVKHRM | 7.763223999 | 7.763223999 | 0 | 0 |
| 59 | GGVSRPT | 7.734213171 | 7.734213171 | 0 | 0 |
| 60 | LQGNHQF | 7.734213171 | 7.734213171 | 0 | 0 |
| 61 | DLMFKGD | 7.704606977 | 7.704606977 | 0 | 0 |
| 62 | QPGSEKY | 7.704606977 | 7.704606977 | 0 | 0 |
| 63 | DQRQKYT | 7.704606977 | 7.704606977 | 0 | 0 |
| 64 | KYQHHPK | 7.704606977 | 7.704606977 | 0 | 0 |
| 65 | DDLITPS | 7.704606977 | 7.704606977 | 0 | 0 |
| 66 | TSKEERL | 7.67438047 | 7.67438047 | 0 | 0 |
| 67 | HEKHKHK | 7.67438047 | 7.67438047 | 0 | 0 |
| 68 | NPYTRPE | 7.67438047 | 7.67438047 | 0 | 0 |
| 69 | AGKQKHI | 7.643507098 | 7.643507098 | 0 | 0 |
| 70 | NATKMTR | 7.643507098 | 7.643507098 | 0 | 0 |
| 71 | QSHSEDY | 7.643507098 | 7.643507098 | 0 | 0 |
| 72 | TNPMSRL | 7.643507098 | 7.643507098 | 0 | 0 |

| Rank | Peptide sequence | subtraction_score | E_cerebrum | E_liver | E_skeletal_muscle |
|---|---|---|---|---|---|
| 73 | HTSPDPM | 7.611958568 | 7.611958568 | 0 | 0 |
| 74 | TVTNGMK | 7.611958568 | 7.611958568 | 0 | 0 |
| 75 | SGLGHLA | 7.579704688 | 7.579704688 | 0 | 0 |
| 76 | DNTVQMK | 7.579704688 | 7.579704688 | 0 | 0 |
| 77 | PAGDNHS | 7.546713197 | 7.546713197 | 0 | 0 |
| 78 | VPASDDK | 7.546713197 | 7.546713197 | 0 | 0 |
| 79 | VNDNNSA | 7.546713197 | 7.546713197 | 0 | 0 |
| 80 | SKVNEKK | 7.546713197 | 7.546713197 | 0 | 0 |
| 81 | RATSLTM | 7.546713197 | 7.546713197 | 0 | 0 |
| 82 | GMHKLTR | 7.546713197 | 7.546713197 | 0 | 0 |
| 83 | VPPFSTN | 7.546713197 | 7.546713197 | 0 | 0 |
| 84 | NTDDAQV | 7.546713197 | 7.546713197 | 0 | 0 |
| 85 | FSSKTTY | 7.512949565 | 7.512949565 | 0 | 0 |
| 86 | SNPRELL | 7.512949565 | 7.512949565 | 0 | 0 |
| 87 | NSKKTEI | 7.512949565 | 7.512949565 | 0 | 0 |
| 88 | TLYKPVD | 7.512949565 | 7.512949565 | 0 | 0 |
| 89 | GHNNTPS | 7.478376784 | 7.478376784 | 0 | 0 |
| 90 | LRTELRL | 7.442955115 | 7.442955115 | 0 | 0 |
| 91 | ISWNPGS | 7.442955115 | 7.442955115 | 0 | 0 |
| 92 | KTAWATR | 7.442955115 | 7.442955115 | 0 | 0 |
| 93 | HHQGGYL | 7.442955115 | 7.442955115 | 0 | 0 |
| 94 | YSSNTMM | 7.442955115 | 7.442955115 | 0 | 0 |
| 95 | IYESATM | 7.442955115 | 7.442955115 | 0 | 0 |
| 96 | RDNGMME | 7.442955115 | 7.442955115 | 0 | 0 |
| 97 | SSSGPPH | 7.442955115 | 7.442955115 | 0 | 0 |
| 98 | NSSFTSL | 7.40664182 | 7.40664182 | 0 | 0 |
| 99 | STLTARS | 7.40664182 | 7.40664182 | 0 | 0 |
| 100 | KLPLTNN | 7.40664182 | 7.40664182 | 0 | 0 |

### Table S4

| motif-defined set | tissue layer | anatomical sample label | motif-matching reads | total reads in anatomical sample label | enrichment ratio | one-sided Fisher’s exact test P value | Benjamini–Hochberg [illegible] value |
|---|---|---|---|---|---|---|---|
| STFHQ_family | parenchyma | dorsal_root_ganglion | 597 | 90485 | 129.942561254 | 0 | 0 |
| STFHQ_family | parenchyma | spleen_red_pulp | 49 | 418872 | 2.30392550209 | 1.24169280774e-07 | 1.36586208852e-06 |
| STFHQ_family | parenchyma | liver_bile_ductal_cells | 46 | 49585 | 18.2709528326 | 1.5712596173e-41 | 2.59257836854e-40 |
| STFHQ_family | parenchyma | appendix_mucosa_epithelial_cells | 32 | 307569 | 2.04909031218 | 0.00015836386507 | 0.00130650188683 |
| STFHQ_family | parenchyma | spleen_white_pulp | 25 | 324804 | 1.51590617493 | 0.0291062826073 | 0.137215332292 |
| STFHQ_family | parenchyma | colon_mucosa_epithelial_cells | 21 | 393551 | 1.05092556482 | 0.439322440641 | 0.99999995911 |
| STFHQ_family | parenchyma | anterior_pituitary_gland | 20 | 197935 | 1.99003668574 | 0.00346166676379 | 0.022847000641 |
| STFHQ_family | parenchyma | liver_around_hepatocytes | 18 | 236058 | 1.50178396942 | 0.0612044955011 | 0.252468543942 |
| STFHQ_family | parenchyma | cerebrum_pvn_neuron | 17 | 186530 | 1.79495643963 | 0.0167704713604 | 0.0922375924823 |
| STFHQ_family | parenchyma | thymus_macrophages | 14 | 207504 | 1.32878661604 | 0.17649366414 | 0.647143435179 |
| STFHQ_family | parenchyma | adrenal_gland_zona_reticularis | 10 | 305988 | 0.643649279371 | 0.947160828732 | 0.99999995911 |
| STFHQ_family | parenchyma | lung_tracheal_epithelial_cells | 10 | 220648 | 0.892593432509 | 0.682514401293 | 0.99999995911 |
| STFHQ_family | parenchyma | thymus_lymphoid_cells | 9 | 277214 | 0.639412367797 | 0.941537769371 | 0.99999995911 |
| STFHQ_family | parenchyma | adrenal_gland_zona_glomerulosa | 8 | 363394 | 0.43357668139 | 0.997988785111 | 0.99999995911 |
| STFHQ_family | parenchyma | pancreas_endocrine_cells | 8 | 271648 | 0.580012238474 | 0.965538516082 | 0.99999995911 |
| STFHQ_family | parenchyma | cerebrum_sn_neuron | 7 | 213900 | 0.644526736734 | 0.916702076068 | 0.99999995911 |
| STFHQ_family | parenchyma | liver_kupffer_cells | 7 | 405347 | 0.340114195954 | 0.999850828482 | 0.99999995911 |
| STFHQ_family | parenchyma | prostate_gland_cells | 6 | 295220 | 0.400275636535 | 0.997339546444 | 0.99999995911 |
| STFHQ_family | parenchyma | skin_adipocytes | 6 | 245312 | 0.48171052952 | 0.985226637554 | 0.99999995911 |
| STFHQ_family | parenchyma | salivary_gland_serous_cells | 5 | 253808 | 0.387988077003 | 0.996117939848 | 0.99999995911 |
| STFHQ_family | parenchyma | testis_interstitial_cells_1 | 5 | 250188 | 0.393601922747 | 0.995564867745 | 0.99999995911 |
| STFHQ_family | parenchyma | adrenal_gland_zona_fasciculata | 4 | 370847 | 0.212431494062 | 0.999992403294 | 0.99999995911 |
| STFHQ_family | parenchyma | cerebellum_medulla_total | 4 | 343742 | 0.229182300325 | 0.999975174652 | 0.99999995911 |
| STFHQ_family | parenchyma | lung_alveolar type 2 cells | 4 | 256161 | 0.307539329869 | 0.999010810592 | 0.99999995911 |
| STFHQ_family | parenchyma | cerebellum_cortex_total | 3 | 441206 | 0.133916326407 | 0.99999995911 | 0.99999995911 |
| STFHQ_family | parenchyma | kidney_tubular epithelial cells | 3 | 212092 | 0.278580458994 | 0.998592367231 | 0.99999995911 |
| STFHQ_family | parenchyma | thymus_dendritic_cells | 3 | 217347 | 0.271844960864 | 0.998875807398 | 0.99999995911 |
| STFHQ_family | parenchyma | prostate_smooth_muscle_cells | 2 | 241580 | 0.163050712556 | 0.999941657249 | 0.99999995911 |
| STFHQ_family | parenchyma | appendix_lymph_nodes | 1 | 249168 | 0.0790426361717 | 0.999997059416 | 0.99999995911 |
| STFHQ_family | parenchyma | cerebellum_4_5cb_medulla | 1 | 72854 | 0.27033375751 | 0.975435700952 | 0.99999995911 |
| STFHQ_family | parenchyma | cerebellum_purkinje_cells | 1 | 317610 | 0.0620096834785 | 0.99999991372 | 0.99999995911 |
| STFHQ_family | parenchyma | prostate_stromal_cells | 1 | 287324 | 0.0685459466304 | 0.999999588126 | 0.99999995911 |
| STFHQ_family | parenchyma | ureter_smooth_muscle_cells | 1 | 59244 | 0.332436965256 | 0.950853428161 | 0.99999995911 |
| STLHQ_family | capillary | thyroid | 15089 | 124306 | 5.14713927207 | 0 | 0 |
| STLHQ_family | parenchyma | spleen_red_pulp | 46268 | 418872 | 4.68378497668 | 0 | 0 |
| STLHQ_family | parenchyma | spleen_white_pulp | 37588 | 324804 | 4.90710557648 | 0 | 0 |
| STLHQ_family | parenchyma | colon_mucosa_epithelial_cells | 29669 | 393551 | 3.19668204009 | 0 | 0 |
| STLHQ_family | parenchyma | thymus_macrophages | 24190 | 207504 | 4.94318595434 | 0 | 0 |
| STLHQ_family | parenchyma | appendix_mucosa_epithelial_cells | 23542 | 307569 | 3.2456250564 | 0 | 0 |
| STLHQ_family | parenchyma | anterior_pituitary_gland | 20549 | 197935 | 4.40215823804 | 0 | 0 |

| motif-defined set | tissue layer | anatomical sample label | motif-matching reads | total reads in anatomical sample label | enrichment ratio | one-sided Fisher’s exact test P value | Benjamini–Hochberg value |
|---|---|---|---|---|---|---|---|
| STLHQ_family | parenchyma | liver_around_hepatocytes | 18296 | 236058 | 3.28651017429 | 0 | 0 |
| STLHQ_family | parenchyma | liver_kupffer_cells | 18146 | 405347 | 1.8982416412 | 0 | 0 |
| STLHQ_family | parenchyma | thymus_lymphoid_cells | 15032 | 277214 | 2.29931864178 | 0 | 0 |
| STLHQ_family | parenchyma | cerebrum_pvn_neuron | 14907 | 186530 | 3.38874674925 | 0 | 0 |
| STLHQ_family | parenchyma | cerebrum_sn_neuron | 13771 | 213900 | 2.72993462215 | 0 | 0 |
| STLHQ_family | parenchyma | adrenal_gland_zona_glomerulosa | 12927 | 363394 | 1.50840356153 | 0 | 0 |
| STLHQ_family | parenchyma | pancreas_endocrine_cells | 11549 | 271648 | 1.80274965943 | 0 | 0 |
| STLHQ_family | parenchyma | adrenal_gland_zona_reticularis | 11271 | 305988 | 1.56190856892 | 0 | 0 |
| STLHQ_family | parenchyma | prostate_gland_cells | 11060 | 295220 | 1.58857200132 | 0 | 0 |
| STLHQ_family | parenchyma | lung_alveolar type 2 cells | 10816 | 256161 | 1.79040474993 | 0 | 0 |
| STLHQ_family | parenchyma | testis_interstitial_cells_1 | 9162 | 250188 | 1.55282088943 | 0 | 0 |
| STLHQ_family | parenchyma | cerebellum_cortex_total | 8999 | 441206 | 0.864869127498 | 1 | 1 |
| STLHQ_family | parenchyma | skin_adipocytes | 8136 | 245312 | 1.40633795995 | 2.42891298731e-194 | 9.83131447245e-194 |
| STLHQ_family | parenchyma | salivary_gland_serous_cells | 7317 | 253808 | 1.22243366307 | 8.47795888014e-65 | 3.00261043672e-64 |
| STLHQ_family | parenchyma | lung_tracheal_epithelial_cells | 6976 | 220648 | 1.34061486123 | 1.45830616489e-125 | 5.63436472797e-125 |
| STLHQ_family | parenchyma | adrenal_gland_zona_fasciculata | 6614 | 370847 | 0.756252760993 | 1 | 1 |
| STLHQ_family | parenchyma | liver_capsule | 6510 | 43588 | 6.33303078411 | 0 | 0 |
| STLHQ_family | parenchyma | liver_bile_ductal_cells | 5918 | 49585 | 5.06083520858 | 0 | 0 |
| STLHQ_family | parenchyma | appendix_lymph_nodes | 5759 | 249168 | 0.980059324539 | 0.941443151094 | 1 |
| STLHQ_family | parenchyma | cerebellum_purkinje_cells | 5754 | 317610 | 0.768198124441 | 1 | 1 |
| STLHQ_family | parenchyma | kidney_tubular epithelial cells | 5676 | 212092 | 1.13479039721 | 1.06324358267e-21 | 3.47598863564e-21 |
| STLHQ_family | parenchyma | prostate_smooth_muscle_cells | 4722 | 241580 | 0.828824458485 | 1 | 1 |
| STLHQ_family | parenchyma | thymus_dendritic_cells | 4531 | 217347 | 0.883970892688 | 1 | 1 |
| STLHQ_family | parenchyma | cerebellum_medulla_total | 3859 | 343742 | 0.476035928239 | 1 | 1 |
| STLHQ_family | parenchyma | prostate_stromal_cells | 3249 | 287324 | 0.479485369623 | 1 | 1 |
| STLHQ_family | parenchyma | cerebellum_4_5cb_medulla | 2557 | 72854 | 1.48824653253 | 4.47333530867e-82 | 1.65318913581e-81 |
| STLHQ_family | parenchyma | liver_hepatocytes | 2182 | 64061 | 1.44430390421 | 6.03029488988e-61 | 2.05030026256e-60 |
| STLHQ_family | parenchyma | ureter_transitional_epithelial_cells | 1128 | 71378 | 0.670104103561 | 1 | 1 |
| STLHQ_family | capillary | testis | 178 | 185308 | 0.0407308418311 | 1 | 1 |
| STLHQ_family | parenchyma | ureter_adventitial_cells | 995 | 62281 | 0.677430984132 | 1 | 1 |
| STLHQ_family | parenchyma | ureter_smooth_muscle_cells | 922 | 59244 | 0.659909074628 | 1 | 1 |
| STLHQ_family | parenchyma | posterior_pituitary_gland | 677 | 60972 | 0.470820951277 | 1 | 1 |
| STLHQ_family | parenchyma | stomach_icc | 561 | 39766 | 0.598202888603 | 1 | 1 |
| STLHQ_family | parenchyma | gallbladder_smooth_muscle_cells | 551 | 55098 | 0.424046337839 | 1 | 1 |
| STLHQ_family | parenchyma | ovary_oocytes | 471 | 105267 | 0.18972572178 | 1 | 1 |
| STLHQ_family | parenchyma | spleen_tubular_endothelial_cells_around_red_pulp | 450 | 47152 | 0.40467832898 | 1 | 1 |
| STLHQ_family | parenchyma | heart_coronary_artery_endothelial_cells | 338 | 87434 | 0.163920739911 | 1 | 1 |
| STLHQ_family | parenchyma | cerebellum_2cb_medulla | 324 | 86354 | 0.159096308787 | 1 | 1 |
| STLHQ_family | capillary | appendix | 51 | 152390 | 0.0141909431352 | 1 | 1 |
| STLHQ_family | parenchyma | ovary_epithelial_cells | 268 | 57162 | 0.198803914343 | 1 | 1 |
| STLHQ_family | capillary | liver | 41 | 1115 | 1.55921693132 | 0.00427679415387 | 0.0134639815955 |
| STLHQ_family | parenchyma | urinary_bladder_smooth_muscle_cells | 129 | 59044 | 0.0926427613787 | 1 | 1 |

| motif-defined set | tissue layer | anatomical sample label | motif-matching reads | total reads in anatomical sample label | enrichment ratio | one-sided Fisher’s exact test P value | Benjamini–Hochberg value |
|---|---|---|---|---|---|---|---|
| STLHQ_family | parenchyma | urinary_bladder_transitional_epithelial_cells | 85 | 29327 | 0.122899138699 | 1 | 1 |
| STLHQ_family | parenchyma | ovary_luteal_cells | 79 | 60892 | 0.0550128830234 | 1 | 1 |
| STLHQ_family | parenchyma | cerebellum_fi_cortex | 68 | 54606 | 0.0528039122534 | 1 | 1 |
| STLHQ_family | parenchyma | prostate_epithelial_cells | 56 | 53699 | 0.0442200655014 | 1 | 1 |
| STLHQ_family | parenchyma | uterus_endometrial_epithelial_cells | 43 | 40431 | 0.0450974021819 | 1 | 1 |
| STLHQ_family | parenchyma | heart_endothelial_cells | 41 | 39437 | 0.0440836493248 | 1 | 1 |
| STLHQ_family | parenchyma | uterus_smooth_muscle_cells | 31 | 41798 | 0.0314487758378 | 1 | 1 |
| STLHQ_family | parenchyma | small_intestine_smooth_muscle_cells | 30 | 88929 | 0.01430458948 | 1 | 1 |
| STLHQ_family | parenchyma | cerebellum_sim_cortex | 25 | 33560 | 0.0315875257715 | 1 | 1 |
| STLHQ_family | parenchyma | cerebellum_3cb_cortex | 21 | 36238 | 0.0245726857583 | 1 | 1 |
| STLHQ_family | parenchyma | cerebellum_4_5cb_cortex | 18 | 39398 | 0.0193729555491 | 1 | 1 |
| STLHQ_family | parenchyma | appendix_icc | 16 | 29554 | 0.0229562669531 | 1 | 1 |
| STLHQ_family | parenchyma | cerebellum_2cb_cortex | 16 | 58380 | 0.0116212660762 | 1 | 1 |
| STLHQ_family | parenchyma | kidney_glomerular_cells | 16 | 180705 | 0.00375445899965 | 1 | 1 |
| STLHQ_family | parenchyma | aorta_smooth_muscle_cells | 8 | 80218 | 0.00422878601767 | 1 | 1 |
| STLHQ_family | parenchyma | spinal_cord_drg_neuron_total | 8 | 100364 | 0.00337994456942 | 1 | 1 |
| STLHQ_family | parenchyma | testis_sertoli_cells | 8 | 44947 | 0.00754721687244 | 1 | 1 |
| STLHQ_family | parenchyma | thyroid_gland_parafollicular_cells | 7 | 44093 | 0.00673171846257 | 1 | 1 |
| STLHQ_family | parenchyma | lung_alveolar type1 cells | 7 | 28649 | 0.010360629068 | 1 | 1 |
| STLHQ_family | parenchyma | colon_smooth_muscle_cells | 6 | 230437 | 0.0011040699522 | 1 | 1 |
| STLHQ_family | parenchyma | lung_club_cells | 6 | 36445 | 0.00698089086498 | 1 | 1 |
| STLHQ_family | parenchyma | skin_sebocytes | 6 | 189865 | 0.00133999719577 | 1 | 1 |
| STLHQ_family | parenchyma | thymus_epithelial_cells | 6 | 43420 | 0.00585947875574 | 1 | 1 |
| STLHQ_family | parenchyma | small_intestine_icc | 5 | 32219 | 0.00658044858557 | 1 | 1 |
| STLHQ_family | parenchyma | kidney_glomerular_mesangial_cells | 4 | 26095 | 0.00649980373186 | 1 | 1 |
| STLHQ_family | parenchyma | thyroid_follicular_cells | 4 | 106547 | 0.00159190196235 | 1 | 1 |
| STLHQ_family | parenchyma | trachea_chondrocyte | 4 | 34063 | 0.00497937287916 | 1 | 1 |
| STLHQ_family | parenchyma | colon_icc | 3 | 62397 | 0.00203870833192 | 1 | 1 |
| STLHQ_family | parenchyma | salivary_gland_duct_cells | 3 | 48203 | 0.00263903250393 | 1 | 1 |
| STLHQ_family | parenchyma | skin_epithelial_cells | 3 | 43666 | 0.00291323418191 | 1 | 1 |
| STLHQ_family | parenchyma | skin_hair_root_cells | 2 | 51374 | 0.00165076087498 | 1 | 1 |
| STLHQ_family | parenchyma | cerebellum_fi_medulla | 1 | 61613 | 0.000688216684721 | 1 | 1 |
| STLHQ_family | parenchyma | cerebellum_sim_medulla | 1 | 71066 | 0.000596672031572 | 1 | 1 |

#### Table S5. Variant counts of the STLHQ motif peptide sets in the capillary and parenchyma datasets across Tier 0–Tier 4 perturbation conditions.

| Tier | Definition | Capillary variants | Parenchyma variants |
|---|---|---|---|
| Tier 0 | ^STLHQ..$ | 5 | 11 |
| Tier 1 | ^ST[IVL]HQ..$ | 6 | 13 |
| Tier 2 | ^ST[TVLI]HQ..$ | 6 | 13 |
| Tier 3 | ^[ST][TL][IVL]HQ..$ | 7 | 14 |
| Tier 4 | ^STL[HNY]Q..$ | 5 | 13 |
